# Safety Recommendations for Temporal Interference Stimulation in the Brain

**DOI:** 10.1101/2022.12.15.520077

**Authors:** Antonino M. Cassarà, Taylor H. Newton, Katie Zhuang, Sabine J. Regel, Peter Achermann, Niels Kuster, Esra Neufeld

## Abstract

Temporal interference stimulation (TIS) is a new form of transcranial electrical stimulation (tES) that has been proposed as a method for targeted, non-invasive stimulation of deep brain structures. While TIS holds promise for a variety of clinical and non-clinical applications, little data is yet available regarding its effects in humans. To inform the design and approval of experiments involving TIS, researchers require quantitative guidance regarding exposure limits and other safety concerns. To this end, we sought to delineate a safe range of exposure parameters (voltages and currents applied via external scalp electrodes) for TIS in humans through comparisons with well-established but related brain stimulation modalities. Specifically, we surveyed the literature for adverse events (AEs) associated with transcranial alternating/direct current stimulation (tACS/tDCS), deep brain stimulation (DBS), and TIS to establish known boundaries for safe operating conditions. Drawing on the biophysical mechanisms associated with the identified AEs, we determined appropriate exposure metrics for each stimulation modality. Using these metrics, we conducted an *in silico* comparison of various exposure scenarios for tACS, DBS, and TIS using multiphysics simulations in an anatomically detailed head model with realistic current strengths. By matching stimulation scenarios in terms of biophysical impact, we inferred the frequency-dependent TIS stimulation parameters that resulted in exposure magnitudes known to be safe for tACS and DBS. Based on the results of our simulations and existing knowledge regarding tES and DBS safety, we propose frequency-dependent thresholds below which TIS voltages and currents are unlikely to pose a risk to humans. Safety-related data from ongoing and future human studies are required to verify and refine the thresholds proposed here.

## Introduction

### Background

Modern neuroscience relies on technologies that modulate neural activity to both illuminate the fundamentals of brain physiology in health and disease, and to treat various neurological and psychiatric conditions. Clinical neuromodulation of the central and peripheral nervous system (CNS, PNS) began in the late 1960s/early 1970s with the use of deep brain stimulation (DBS) and spinal cord stimulation for chronic pain^1^. In contrast to implanted electrodes, early non-invasive systems for transcranial electrical stimulation (tES) used high intensity currents to directly affect brain activity via scalp electrodes, e.g., electroconvulsive therapy (ECT)^2^. Today, more sophisticated invasive and non-invasive stimulation technologies are available to treat several neurological and psychiatric diseases^3^, or to study cognitive functions^4,5^.

DBS is a system for delivering electrical currents directly into the brain via an invasive surgical procedure wherein electrodes are implanted in disease-specific subcortical structures (e.g., globus pallidus (GP), subthalamic nucleus (STN)). Currents are applied through these electrodes to target the pathophysiology of neural circuits associated with various disease states; the treatment of Parkinson’s disease (PD) in particular has benefited from this approach^6^. DBS first garnered public attention in 1987, when Benabid et al. reported successful tremor suppression in PD patients by applying high frequency stimulation to the thalamus^7^. Since then, it has also been investigated as a treatment for other conditions, including obsessive compulsive disorder (OCD), major depression (MD) and Alzheimer’s disease (AD)^6^. However, the invasiveness of the procedure and its associated risks are considerable, relegating DBS to a category of “last resort” options for chronic refractory disease where patients fail to respond to other treatments.

Non-invasive brain stimulation (NIBS) methods, such as tES and transcranial magnetic stimulation (TMS) are often used in healthy participants^4^ but have also been the focus of clinical investigations^8^. In 2008, TMS was approved by the U.S. Food and Drug Administration (FDA) for patients with treatment-resistant depression^9^. However, tES and TMS have the limitation that they mainly target the cortex, in the case of tES by delivering currents through scalp electrodes, and in the case of TMS by inducing currents in the head using an electromagnetic (EM) coil, with the electrodes or coil being positioned over a specific target area. Neither technique is well-suited for direct modulation of deep brain structures, since the intensity of stimulation that would be required would tend to activate overlying cortical areas, resulting in unwanted side effects, off-target stimulation, and the stimulation of peripheral nerves^10^.

Recently, temporal interference stimulation (TIS), a new form of tES, has been proposed as a method to enable targeted, yet non-invasive stimulation of structures located deep inside the brain^11^. The method uses two or more harmonic (sinusoidally varying) electric (E-)fields that are applied through scalp electrodes at slightly different frequencies within the kHz range (e.g., 1.01 kHz and 1 kHz). While these frequencies are themselves too high to induce neural firing, the difference frequency between the fields, which causes modulation of the field “envelope” at that frequency (i.e., at 10 Hz in the example above; see also Figure 1), is within the physiological range of electrical brain activity (1–100 Hz) and is able to modulate neural activity. This means that brain regions or networks can be electrically activated at targeted locations within the brain without necessarily driving neighboring or overlying regions. Furthermore, by varying the magnitudes of the electrical currents applied to each of the electrode pairs, a certain degree of electronic focus steering is possible without moving the stimulation electrodes.

**Figure 1.**
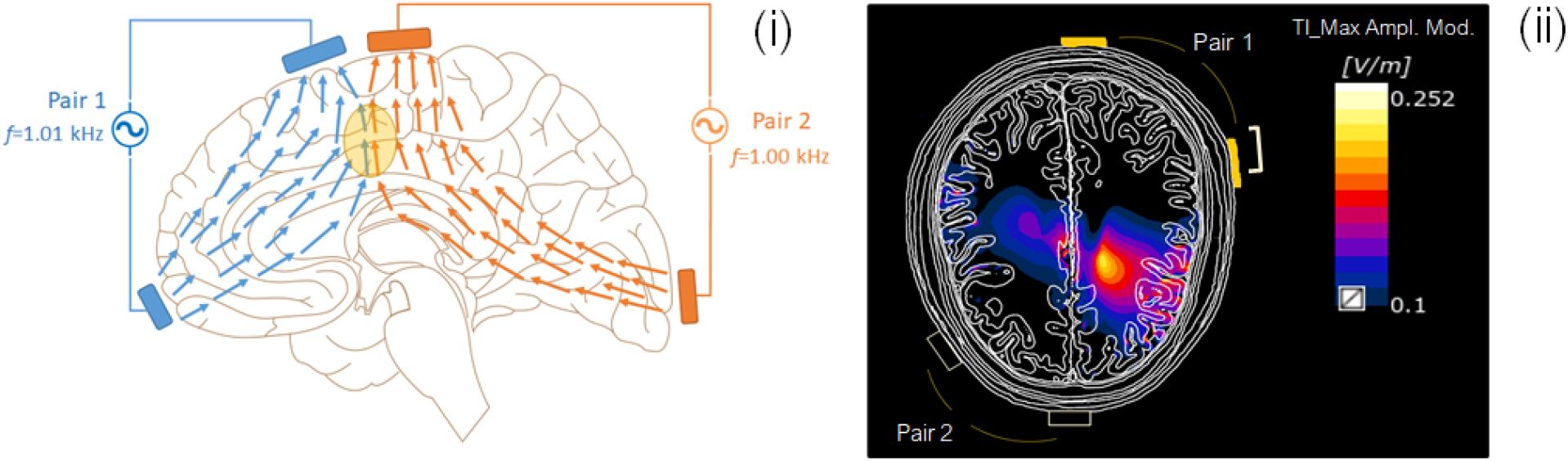
(i) Schematic representation of the TIS concept. Two currents in the kilohertz range (1.01 kHz, in blue; 1 kHz, in orange) are simultaneously applied to the head via two pairs of scalp electrodes. The applied currents generate E-fields inside the brain (colored arrows). The envelope of the total field varies with time at the difference frequency (here, 10 Hz), which is capable of modulating neural activity. The location of neural modulation is spatially constrained to the region where the currents overlap (yellow region), producing the strongest field interference effect. (ii) Simulated modulation envelope magnitude map from the setup described in *<u>Computational Characterization and Comparison of Electric Brain Stimulation Modalities</u>*.

### Motivation

TIS holds promise for a variety of clinical and non-clinical applications^12–14^. Like other tES technologies, it alters neuronal excitability at the subthreshold level. However, TIS differs from technologies such as transcranial alternating current stimulation (tACS) and transcranial direct current stimulation (tDCS) in the following ways:

- Neural activity in the stimulated brain region is modulated at the difference frequency (i.e., the frequency difference between two kHz range applied currents), which is chosen to be within the physiological range (e.g., 10 Hz in Figure 1). By contrast, tACS and tDCS exert their effects directly at the applied frequency (tACS), or in the absence of frequency content (tDCS).
- TIS requires a more complex electrode configuration than either tACS or tDCS, involving at least two pairs of electrodes. As the number of electrode pairs grows, increasingly complex E-field modulation patterns can be achieved. The complexity of such patterns is further exacerbated by the impacts of head anatomy and tissue heterogeneity. Thus, optimal stimulation sites must be carefully planned through modeling and simulation.
- TIS uses multiple current intensities, one per electrode pair, which must be precisely calibrated to allow for steering of the stimulation site focus. This choice simultaneously dictates the intensity of stimulation and the distribution of TI modulation for a given electrode configuration.

The recency of TIS as a brain stimulation modality means that published data regarding its effects in humans (adverse or therapeutic) are sparse. Therefore, it is important to examine whether TIS may pose different risks to humans as compared to better-established tES technologies such as tACS and tDCS. To this end, we surveyed scientific literature and public databases for the known risks and adverse events (AEs) associated with related brain stimulation technologies. Here, we define an AE as any undesirable experience associated with the use of a stimulation device in a person. While TIS differs in substantial ways from other forms of tES (e.g., carrier frequency, modulation, as highlighted above), we attempt to integrate existing empirical knowledge regarding tES safety with (bio)physical principles to establish boundaries for the safe use of TIS in research and clinical settings. A detailed view of the landscape of TIS safety will require the emergence of a more substantial body of literature concerning its use in practice.

We selected tACS and tDCS for inclusion in this paper, as they are well-studied methods for subthreshold cortical electrical stimulation with key similarities to TIS, namely their mechanisms of action (electrical), their typical stimulation intensities (mA), and their mode of current delivery (surface scalp electrodes). Like TIS, tACS uses alternating current for stimulation, and is thus of particular relevance. Unlike tACS and tDCS however, TIS can be used to apply localized stimulation to deep structures in the brain. At the same time, it may create unwanted off-target stimulation due to the interferential nature of the stimulation itself and the complex anatomical and dielectric heterogeneity and anisotropy of the brain. Like TIS, DBS stimulates deep brain regions locally. Therefore, although DBS is an invasive procedure and electrode design and stimulation parameters are quite different from TIS, DBS literature was also considered as it may provide useful information concerning AEs that result from (i) direct, focal stimulation of deep brain structures; and (ii) strong fields in off-target, nearby regions (r ≥15 mm from the active electrode). TMS and related techniques were not considered in this document, as their stimulation mechanism (suprathreshold magnetic pulses) is not readily comparable to that of TIS.

We extrapolate these data to the case of TIS considering the (bio)physical principles that are thought to support its action in the brain and provide a rationale for which of the reported AEs may or may not be applicable to TIS, taking into account the different delivery methods, frequencies, and specificity of modulation. This work, therefore, is a first step towards delineating the parameter space of safe and effective TIS in humans, with an eye towards future trials and experiments that will help close existing gaps in the literature.

### Biophysics of Electrical Stimulation and Metrics of Exposure

Therapeutic interventions in the CNS generally target the modulation of neural spiking in a manner consistent with restoration or enhancement of healthy activity patterns, or suppression of pathological activity. While such interventions may exert their effects via electric, magnetic, mechanical, thermal, chemical, or other means, in the context of TIS safety, we restrict ourselves to the consideration of neuromodulation achieved via applied E-fields. In general, electrical stimulation may act on either the sub- or suprathreshold activity of neurons. In the subthreshold regime, neuromodulation alters membrane polarization and synaptic efficacy, while for suprathreshold stimulation, APs are induced directly. Moreover, the effects of electrical stimulation may be considered with respect to their immediate effects, or their long-term influence on network plasticity.

We present this information here to facilitate comparisons between TIS and other, better-established modalities of electrical neurostimulation in terms of the underlying biophysical mechanisms. In this way, we aim to inform a broader discussion of TIS safety, ultimately leading to safer and more effective stimulation paradigms. *Appendix B* provides a review of the relevant exposure and interaction biophysics, beginning with an overview of the physical interactions between E-fields and neurons. In addition, we discuss the resulting impacts on neural physiology in the sub- and suprathreshold regimes, distinguishing short- and long-term effects. While our focus is the electrophysiological consequences of electrical neurostimulation, other aspects, including heating, charge accumulation, electrochemical reactions, and electrode-related effects, are also considered in view of potential AEs.

Based on our review of neuromodulation biophysics, we identified several exposure metrics with particular relevance for safety, namely: E-field magnitude and heterogeneity in the brain and skin, total current magnitude and current density, temperature increase, and injected charge per phase. In *<u>Computational Characterization and Comparison of Electric Brain Stimulation Modalities</u>* we compare neurostimulation modalities in terms of these quantities.

### Transcranial Electrical Stimulation (tDCS and tACS)

#### Basic Principles

The category of tES encompasses several stimulation protocols, including but not limited to tACS and tDCS. To administer tES, current is passed through two or more electrodes placed on the surface of the scalp via a battery-powered device^15^. The current is transferred through the scalp and skull into the brain, where it induces subthreshold changes in the transmembrane potential of cortical neurons. These changes render the affected neurons either more or less excitable, depending on the polarity of the stimulation. Here we focus on the most common stimulation protocols, parameters, and montages (i.e., electrode sizes and placements) reported in the literature^16,17^. In typical applications, tES stimulation amplitudes fall between 1–2 mA (peak to peak for tACS) and do not exceed 4 mA. In tDCS, constant, direct current is passed through the electrode, while tACS makes use of alternating currents. Electrodes used for tES are generally rectangular or circular with coverage areas between 1–50 cm^2 17^. The particular combination of current amplitude and electrode size determines the current density delivered at the electrode surface. For example, high-density tES (HD-tES) is an emerging technology that seeks to provide targeted stimulation, in part by employing small electrodes. HD-tES thus generates higher current densities than traditional tES for the same amplitude of delivered current^17^. Similarly, the amplitude of current delivered and the treatment duration determine the total charge dosage. Both total charge and charge density are factors that critically influence the safety and effectiveness of stimulation.

#### tDCS

Most commonly, tDCS is applied via a single anode-cathode pair of electrodes. By convention, this entails that positive electrical charge flows from anode to cathode, such that the anode accumulates negative charge, and the cathode positive charge. Consequently, the first-order effects of tDCS depend on either depolarization at the site of the anodic electrode, or hyperpolarization around the site of the cathodic electrode. Typically, experimental protocols aim to either increase neuronal excitability by placing an anodic electrode over the target, or decrease excitability with a cathodic electrode^15^. Note that while anodic (cathodic) stimulation is excitatory (inhibitory) for bundles of pyramidal cells oriented with their principle axes orthogonal to the electrode surface, the scenario may be reversed for different orientations, for example within cortical gyrifications, or different neuron types/axons^18^. Additional factors that may influence outcome include stimulation intensity and duration, coincident pharmacological interventions^19^, inter-subject variability^15^, brain state^16^, and morphology-dependent neural physiology^20^. Furthermore, computational modeling studies have shown that most of the current does not enter the cortex at all, but is either concentrated on the scalp just below the electrodes, or shunted through the comparatively low-impedance cerebral spinal fluid (CSF) surrounding the cortex^21,22^. During operation, current is generally ramped up over a period of seconds to the desired intensity level, and ramped down at the end of the stimulation period. Stimulation is applied for a duration lasting from a few seconds up to one hour within a single session, though most commonly in the range of 10–30 minutes with a total delivered charge ≤7.2 coulombs^16^. The working principles of tDCS have yet to be fully clarified^23^. For details concerning various neurophysiological mechanisms that have been hypothesized to underlie the effects of tDCS, we refer the reader to several reviews dedicated to this subject^24–26^.

#### tACS

Similar to tDCS, tACS is applied via two or more electrodes placed on the surface of the scalp. One “active” electrode is placed over the brain region of interest and one or more “reference” electrodes are located in a region presumed not to interact with the experimental paradigm. As with tDCS, current flows between the active and reference electrodes, and brain regions that lie in the path of current flow are most likely to be stimulated^27^. Unlike tDCS, however, the primary physical mechanism is not the depolarization or hyperpolarization of a cortical region, but rather entrainment of brain oscillations at the stimulation frequency^28^. The resulting physiological effects vary considerably with the applied frequency^29^. A recent review analyzed 57 studies using currents <2.3 mA and frequencies between the theta and gamma bands of oscillatory activity, examining frequency-specific short- and long-term effects on various cognitive functions^4^. The results of the review suggest that stimulation may affect attention (alpha: 8–13 Hz, beta: 13–30 Hz), intelligence (gamma: 30–80 Hz), visual and auditory perception (alpha and gamma, respectively), executive function (beta), decision-making (beta), motor learning (alpha and gamma), working memory (beta) and declarative memory (theta: 4–7 Hz and gamma). Importantly, the authors provide information regarding electrode montages, current intensities and frequencies, and offer a summary of the impacts of tACS categorized by brain function. The studies reviewed, however, were conducted mainly in controlled laboratory environments, with uncertain translation to real-world settings^4^. In addition, as in tDCS, the underlying brain state may alter the effects of stimulation^4,16^.

#### tES and AEs

As of publication, neither tACS nor tDCS have received FDA approval, curtailing their adoption as clinical therapies. Thus, data on AEs in humans is confined primarily to ad hoc reports in academic studies and clinical trials. Additionally, the breadth and depth of available information concerning AEs is heavily skewed towards tDCS (versus tACS) in the tES literature, biasing the results of this survey.

The most frequently reported AE for tES is skin irritation (including tingling, burning, itching, and erythema), closely followed by fatigue, headaches, and phosphene induction^16,17,30^. Reporting of these AEs, however, is neither systematic nor universal in tES studies. In fact, multiple reviews and meta-analyses have found that AE reporting for tES is often either inadequate or entirely absent. Among 158 sham-controlled studies surveyed in a review of tDCS safety, 43 made no mention of AEs, while 42 provided only cursory AE reports^31^. Similarly, Brunoni et al. found that among 172 primary research articles on tDCS (encompassing 209 studies), 92 did not report the occurrence or absence of AEs^32^. Therefore, the tES literature is likely biased towards an underrepresentation of AEs.

#### Prevalence of Specific AEs

In a review of the tDCS literature, Brunoni et al. attempt to quantify the prevalence of common AEs by counting the number of mentions per category. Among 117 studies that reported on AEs, 74 referred to the occurrence of at least one AE, while 43 reported that no AEs were experienced by subjects. In order of decreasing prevalence, itching (39.3%), tingling (22.2%), headache (14.8%), general discomfort (10.4%) and burning sensations (8.7%) were reported for at least one subject during active stimulation^32^. The frequency of each AE was also compared against a sham stimulation condition, in which current was applied in a short ramp-up phase to reach a target amplitude, and subsequently switched off. The prevalence of AEs during the sham stimulation were: itching (32.9%), tingling (18.3%), headache (16.2%), general discomfort (13.4%) and burning sensations (10%), reported as a percentage of the 117 studies in which at least one subject experienced an AE. Hence, the authors note that the prevalence of AEs is similar between active and sham conditions.

Another important metric is the *number* of subjects within a study that experience a particular AE. In a large-scale tDCS meta-study, Kessler et al. subjected a cohort of 131 healthy volunteers to 277 tDCS sessions across eight individual studies to compare the prevalence of AEs during and after stimulation^33^. Electrodes were either 5x5 cm or 5x7 cm, and the applied current amplitude was 1.5 mA in each study. Real and sham stimulation sessions lasted between 10 and 20 min. Similar to the results reported by Brunoni et al., it was found that the most common AE during stimulation was skin irritation, specifically tingling sensations, reported by 76.9% of subjects across the 277 sessions. Other common AEs included itching (68.2%), burning sensations (54.2%), difficulty concentrating (35.7%), pain (24.9%) and fatigue (20.9%). Similarly, AE prevalence immediately following stimulation was dominated by tingling and itching sensations, though such after-effects were reported in fewer sessions: 24.9% and 25.6% for tingling and itching, respectively. It was concluded that while AEs are frequent in tDCS studies, they are benign, and do not pose a medical hazard^33^.

#### Moderate to Severe AEs

Several large-scale reviews address reports of AEs during tES whose severity exceeds skin irritation and general discomfort. Most notable among these are persistent skin lesions and mania/hypomania^17,30,34^. See Table A.1 in *Appendix A* for a summary of moderate to severe AE occurrence across tES studies.

Skin lesions have been reported in multiple studies during the application of tDCS with current amplitudes <2 mA^17,30,34^. Lesions were described alternatively as brown, crusty ulcerations beneath the electrodes or blister-like atrophic scars. Multiple mechanisms have been proposed to account for the appearance of lesions, including: tissue burning following desiccation of the sponge electrolyte^35^, toxic reactions to tap water constituents or impurities^34^, toxic electrochemical reaction products^36^, and pH changes in the skin^37^. In some cases, skin burns were reported as a result of improper electrode placement^38^. Data from at least one clinical trial indicates that the risk of skin lesions is best predicted by the contact medium, and not the site of application, the phase (anodic vs. cathodic), or the level of discomfort experienced by subjects^34^. Lesions that occurred using tap water were linked to chemical skin damage by alkaline hydrolysis due to high local concentrations of calcium carbonate^34^. Lesions below the cathode may be due to direct current iontophoresis causing alkaline accumulation under the negatively charged electrode^39^. Changes in the constitution of the gel to a black paste or white powder have also been described^40^. However, the risk of skin lesions or burns is negligible when adhering to standardized tDCS protocols^41^.

Hypomania and mania are mood disorders characterized by unusually high energy, elatedness, or irritability, and are frequently comorbid with bipolar disorder^42^. Importantly, in studies where hypomania/mania was reported, subject populations were frequently depressive or bipolar, and therefore may have been administered pharmacological interventions concurrently with tDCS. Hence, Antal et al. urge caution when prescribing tES therapies to such populations and recommend that patients be screened regarding their propensity for manic episodes^17^.

To date, there is little evidence of seizure induction by tES, even in epileptic populations^17^. Exceptionally, a single case study reported the occurrence of seizure in a child with idiopathic infantile spasm and spastic tetraparesis who had been seizure-free for one year prior to tDCS treatment^43^. In this case, a partial onset seizure was observed four hours after application of tDCS to the right motor cortex for 20 minutes. However, it is not clear whether tDCS caused the seizure, and it is the only report of seizure associated with tES in the literature to date.

#### AEs and Stimulation Parameters

Various stimulation parameters may affect the risk of AEs. In a recent meta-analysis of the literature, Nikolin et al. investigate the relationship between cumulative charge and AE incidence^31^. Cumulative charge summarizes tDCS exposure as a function of stimulation amplitude, stimulation duration, and number of sessions. Among the studies surveyed, stimulation amplitudes, durations and number of sessions spanned 1–2.25 mA, 2–40 min, and 3–30 sessions, respectively^31^. However, AE incidence was not found to vary significantly with cumulative charge. Hence, the authors conclude that within the range of stimulation parameters commonly used in tDCS experiments, increasing the number of sessions, amplitudes and/or session durations does not increase the probability of AEs. An important limitation of many tDCS studies is the short duration of follow up, leaving long-term effects (order of months) insufficiently characterized^44^. The therapeutic effects of some tES applications act primarily through the potentiation of neuroplasticity over many repeated sessions, underscoring the importance of describing the incidence of long-term AEs.

With respect to tACS, far fewer studies and reviews have been conducted regarding safety, partly due to its comparatively recent emergence as a neurostimulation modality^27^. Compared to tDCS, tACS has been reported to produce milder skin-related AEs, with further attenuation at higher frequencies^30^. In general, tACS-related AEs vary considerably with the applied frequency. For example, tACS in the 8–40 Hz range (particularly 10–20 Hz) is known to induce phosphenes when the active electrode is situated sufficiently near the eye^17,30^. Dizziness and headache have also been reported for various electrode montages and stimulation frequencies^17,30^. Some studies suggest that posterior electrode montages are more likely to elicit dizziness, possibly as a result of proximity to the vestibular nerve^30^. Common targets of tACS include the motor cortical areas, cerebellum, and visual system via transorbital stimulation. In one high-frequency tACS study (n.b. the similarity to TIS carrier frequencies), researchers applied 1 mA of current at 5 kHz over the primary motor cortex (4x4 cm active electrode, 7.5x6 cm return electrode) for 10 min^45^. The authors report that 70.6% and 35.6% of the 18 subjects experienced mild tingling and fatigue, respectively. At the extreme end of applied current amplitude, Antal et al. report on one instance of the theta-burst protocol^46^ applied at 10 mA, wherein bursts of 5 kHz alternating current were delivered for either 1 ms or 5 ms through an active electrode located over the primary motor cortex^17^. In this condition, four of 14 subjects withdrew from the experiment due to painful skin sensations. Of the remaining 10 volunteers, five subjects reported mild to moderate skin tingling.

Collectively, these results suggest that electrode interactions with the skin are the most commonly reported AEs in tDCS studies for typical stimulation amplitudes (<4 mA, ≤30 min stimulation) and electrode montages (5x5 cm – 5x7 cm electrodes). For low current amplitude tES stimulation (<4 mA) increasing stimulation intensity and total accumulated charge (e.g., via repeated sessions) does not appear to increase the occurrence of AEs. However, these conclusions are limited to short-duration treatment/experiment paradigms (30 sessions maximum).

Moderate to severe tES-related AEs, while concerning, are rare for typical stimulation parameters and electrode montages. Bipolar and depressive patients may be at increased risk for hypomanic or manic AEs, additionally subject to the influence of concurrent pharmacological interventions. Patients should be screened for the use of certain medications, and any history of mood disturbances. Furthermore, proper skin and electrode preparation is necessary to mitigate the risk of persistent skin lesions, and subjects should be encouraged to report any intra-stimulation discomfort to prevent further skin irritation^17^. Despite one report of seizure, many thousands of sessions of tES have been performed without incident. Thus, patient history and seizure propensity should be considered prior to tES treatments, but do not necessarily constitute strict exclusion criteria. Overall, for the electrode montages and stimulation parameters reported in tACS literature, there are no documented cases of serious AEs.

### Deep Brain Stimulation

#### Basic Principles

DBS is an invasive procedure in which pulses of electric current delivered via surgically implanted electrodes are used to modulate the activity of deep brain regions for the treatment of various neurological and psychiatric disorders. The electrodes are implanted stereotactically at disease-specific target regions and connected to a programmable implanted pulse generator (IPG), typically located below the clavicle^47^. The IPG contains a rechargeable or single-use battery, electronic components to generate the stimulus waveform(s), and output cables that connect to the implanted electrodes^47,48^. Commonly, DBS employs a monopolar or bipolar electrode configuration; in the former case, the contact serves as the (negative) cathode and the IPG as the (positive) anode, resulting in a relatively wide spread of current. In the latter, both contacts serve as electric poles, resulting in more spatially confined currents^49^.

A standard DBS electrode has four equally spaced contacts (common dimensions: 1.5 mm diameter, 1.5 mm contact length, 0.5 mm intercontact spacing) but designs vary with regard to the number, shape and spacing of the contacts^50,51^. During programming, the active contacts may be altered, allowing for fine-tuning of the stimulus location. Recently, there have been several important advances in DBS technology, including the development of “directional” electrodes which enable more precise control over the volume of stimulated neural tissue^47,51^. Furthermore, directional DBS may avoid the need for complex stimulation protocols such as bipolar stimulation, frequency/pulse-width modulation, or interleaving^50,52^.

In typical DBS applications, current pulses are applied through selected contacts at a pre-programmed frequency (generally >100 Hz), pulse amplitude (current or voltage), and pulse width. For example, patients receiving STN stimulation for PD may be treated with currents in the range of 0.8–2 mA at 130 Hz, and a pulse width of 60 μs^53^. However, these parameters must be optimized for each patient depending on the disease, brain target, and individual differences in anatomy and physiology^47^. In recent years, a trend towards multi-target stimulation has further ramified the complexity of treatment protocols, increasing the importance of pre-operative planning and simulation^54^. Stimulation is usually administered chronically over a duration of weeks, months or years and can be controlled by patients or clinicians via external devices with a radiofrequency (RF) or bluetooth connection. Some modern devices are also controllable through dedicated software on smartphones or tablets. While manual adjustment of DBS parameters is the standard of care as of publication, the feasibility of “adaptive” or “closed-loop” DBS, in which the delivery of stimulation is adjusted dynamically in response to certain biomarkers^55^, has recently been demonstrated in patients with PD^56^. Krauss et al.^50^ and Harmsen et al.^54^ provide comprehensive overviews of technical developments in DBS systems, and advances in stimulation waveform/pattern design, respectively.

Despite the well-established empirical success of DBS, its underlying physiological mechanisms are not yet fully understood. Prominent theories implicate the disruption of pathological brain circuits via the modulation of physiology across scales spanning ion channels, protein expression, single cell dynamics, and networks^6,57^. Regardless of the precise mechanism, it is apparent that high-frequency stimulation tends to reduce neural activity locally, effectively creating a reversible lesion^6,47^. A full description of the mechanisms of action will need to account for variations in physiology among target locations, and chronic adaptations that occur with prolonged exposure^6^. For example, withdrawal-like symptoms have been documented in three PD patients following the cessation of DBS stimulation (duration of eight years) in the STN, indicating long-term changes in the basal ganglia loop^58^. Thus, DBS likely exerts both local and network-level electrical and neurochemical effects by modulating excitability, oscillatory activity, synaptic plasticity, and possibly also by potentiating neurogenesis^48^.

### DBS and AEs

DBS has a broad spectrum of potential therapeutic indications of which the FDA has recognized the following five, as of publication: essential tremor (ET) and PD-associated tremor^59^, PD^60^, drug-resistant epilepsy^61^, dystonia^62^, and OCD^63^, with the last two carrying a humanitarian device exemption^64,65^. Brain targets for each pathology may vary, but include, respectively: the ventral intermediate nucleus (VIM), the STN or globus pallidus internus (GPi), the anterior nucleus of the thalamus (ANT), the STN and GPi, and the anterior limb of the internal capsule (ALIC)^64,65^. In addition, DBS is presently being evaluated in human clinical trials worldwide for a variety of other disorders and brain targets^54^.

In general, achieving maximal clinical benefit for each patient requires personalizing the stimulation configuration through careful selection and fine-tuning of active electrode contacts, frequencies, pulse widths, and voltages^66^. Nevertheless, placement errors on the order of millimeters are commonplace, and may result in AEs and/or reduced treatment efficacy. Various programming strategies exist to mitigate AEs caused by the spread of current into brain structures adjacent to the target region^67^. For example, interleaved programming^68^ and the use of directional electrodes^52^ have been proposed to minimize stimulation-induced AEs and maximize clinical benefits. Table A.2 in *Appendix A* provides a list of commonly reported DBS targets across the main clinical indications, Table A.3 gives an overview of the range of reported DBS stimulation parameters, and Table A.4 provides a list of reported stimulation-related AEs. Note that due to experimental diversity and limited AE reporting within the surveyed DBS literature, it was not feasible to establish a direct link between specific stimulation parameters, target structures, and reported AEs.

The appreciable variation in DBS brain targets across diseases is apparent in Table A.2, with stimulation of the GPi, STN and various thalamic nuclei reported most frequently for movement and psychiatric disorders. Typical stimulation parameters span frequencies up to 190 Hz, pulse widths up to 450 μs, and voltage and current intensities up to 14 V and 8 mA, respectively. However, the optimal combination of parameters varies according to the clinical effect desired^69^. Stimulation frequencies for chronic pain relief tend towards the lower range in accordance with the supposition that DBS of the thalamus or periaqueductal grey matter (PAG) at lower frequencies (<50 Hz) causes analgesia, whereas higher frequency stimulation (>70 Hz) results in hyperalgesia^57^. Stimulation-induced AEs reported in the DBS literature are diverse, including abnormal taste sensations, tingling of the scalp, headache, confusion, gait disruption, balance or speech disturbances, depression, psychosis and seizures (Table A.4). However, apart from those AEs that resolve with the adjustment of parameter settings, it is difficult to establish an irrefutable link between a given AE and stimulation; this is compounded by the lack of consistent associations between stimulation parameters, brain target, and AEs. A recent review investigated the relationship between stimulation parameters, treatment outcomes, and AEs for STN stimulation in PD patients^70^. Notably, the authors observed an overlap of parameter values for positive clinical effects and AEs, with a tendency towards AEs at higher intensities (threshold for AEs: 1.3–3.4 mA and 2.04 V). With respect to frequency, results revealed that beneficial effects were present between 50–185 Hz and that the therapeutic window decreased with increasing pulse width.

The information provided in Tables A.2–A.4 has several limitations. First, we note the possibility of selection bias, since literature with negative findings was generally not considered (and may be less likely to be published). Furthermore, we expect a bias towards underreporting of AEs since most studies focused on other endpoints, and also towards overrepresentation of specific studies and patients given the overlap between reviews. Standardized procedures and best practices for the collection and reporting of AEs are lacking and likely contribute to uncertainty surrounding AEs in the literature^71^. In addition, DBS is often administered concomitantly with medication (e.g., Levodopa for PD) so that distinguishing stimulation-induced AEs from the effects of pharmacological treatments, disease progression, or comorbidities is difficult^72^. Studies involving DBS are often observational and unblinded, are restricted to small sample sizes or single patients, relate to highly patient-specific diseases, symptoms, and anatomy, involve a diverse range of target structures and stimulation parameters, and employ variable follow-up periods. Furthermore, randomized controlled trials and systematically collected data over longer durations are lacking for most DBS indications. Finally, the descriptions of the brain targets themselves may be unclear due to inconsistent or ambiguous nomenclature across studies or even within the same study^73^.

### AEs from Regulatory and Clinical Databases

#### PMA

The PMA records for three FDA-approved DBS implants (Brio Neurostimulation System, Abbott Medical, PMA Nr. P140009, Vercise DBS, Boston Scientific, PMA Nr. P150031, Activa DBS System, Medtronic, PMA Nr. P960009) indicate that the most common AEs related to electrical stimulation are: anxiety, confusion, depression, disequilibrium, dyskinesia, dystonia, edema, hallucinations, headache, nausea, psychiatric changes/disturbances, seizure or convulsion, sleep disturbances, tremor, and visual disturbances.

#### PMS

AEs related to electrical stimulation reported in the PMS data for the categories *Nervous System, Mental Emotional and Behavioral Disorders, Musculoskeletal System*, and *Skin and Subcutaneous Tissue* for product code MHY (Stimulator Electrical, Implanted, for Parkinson Tremor), NHL (Stimulator, Electrical, Implanted, for Parkinsonian Symptoms) and OLM (Deep Brain Stimulator for OCD) included: anxiety, burning sensations, cognitive changes, confusion, dementia, discomfort, disorientation, dizziness, dyskinesia, dysphasia, dyspnea, electric shock, emotional changes, fall, fatigue, irritability, lethargy, loss of consciousness, malaise, memory loss/impairment, muscle spasms, muscular rigidity, neurologic deficiency/dysfunction, numbness, pain, paresis, seizures, shaking/tremors, sleep disturbances, tingling, twitching, and weakness.

### Temporal Interference Stimulation

#### Basic Principles

TIS offers significant advantages over traditional tACS, including focal steerability, and localized stimulation of deep brain target regions without recruitment of overlying structures^11^. Furthermore, the high frequency carrier signals (typically 1–5 kHz, with difference frequencies ranging between 5–50 Hz) result in significantly reduced cutaneous discomfort/sensation, which is the primary limitation on current density in conventional tACS and tDCS (current density <0.1 mA/cm^2^)^30^. During TIS, electrode currents of 1–2 mA per channel at 2 kHz have been applied without or with minimal sensation (personal correspondence with collaborators).

TIS is distinguished from conventional tACS in that the frequency of applied currents is too high (>1 kHz) to induce neuromodulation at typical current strengths due to the intrinsic low-pass filtering properties of neural membranes^74^. Instead, two or more pairs of electrodes are simultaneously active, but with small differences in the frequency of the applied currents, resulting in low-frequency temporal modulation of the signal envelope at the difference frequency of the high-frequency carriers. Empirically, it has been observed that neurons are able to “demodulate” the low-frequency field modulation without responding to the high-frequency carrier fields. It is important to note that the frequency spectrum of the exposure only contains the carrier frequencies; thus, the superposition of pure high-frequency fields lacks any low-frequency content. Therefore, in their passive response, neural membranes should theoretically attenuate both amplitude-modulated and unmodulated oscillations equally, abolishing the excitatory effects of TIS. In spite of this, mouse hippocampal neurons exhibit near identical responses to TIS as to conventional stimulation at the TI difference frequency^11^. Thus, the active, electrophysiological response must be responsible for the demodulation. In *Appendix B*, we review the current state of knowledge regarding the biophysical mechanisms giving rise to neuromodulation by temporally interfering electrical fields applied to the brain. Despite its promise, TIS has so far eluded efforts to definitively explain the mechanism by which neurons are able to demodulate envelope fluctuations. Since the present understanding of such mechanisms is incomplete, we are careful to delineate what is known from areas where consensus has yet to emerge, with an eye towards promising avenues of ongoing research.

### Mechanistic Hypotheses and Relevant Dosimetric Quantity

Regarding the observed capacity of neurons to demodulate TI field oscillations, unpublished hypotheses voiced in discussions with researchers include: (i) nonlinear frequency mixing in the subthreshold regime (possibly caused by nonlinear membrane capacitances), (ii) periodic high-frequency conduction blocking (CB) (iii) network amplification of synchronized perturbations, (iv) stimulation resulting from temporal field amplitude variations, and (v) localized ion-channel effects (e.g., related to ion diffusion). However, these theories conflict to varying degrees with empirical observations. For a more comprehensive discussion of prevailing theories and published mechanistic investigations, see *Appendix B*.

As a consensus understanding of the biophysical mechanisms underlying TIS is lacking, the results of modeling efforts in the design of safe and efficacious protocols, treatment plans (electrode placement, carrier and modulation frequencies, temporal stimulation adaptation, brain region targeting) and stimulation devices must be interpreted with caution. This also complicates the identification of an exposure metric that can be used for treatment optimization and safety assessments. Based on the observation that stimulation can be “steered” by changing the current ratio(s) without moving the electrodes, and that the stimulation focus shifts towards the electrodes providing the *weaker* current^11^, the amplitude of the modulation envelope has been proposed as a suitable TIS exposure metric. The spatial distribution of its projection along a unit vector *<u>n</u>* (e.g., parallel to a principal neuronal orientation) for a two current configuration may be calculated as follows:

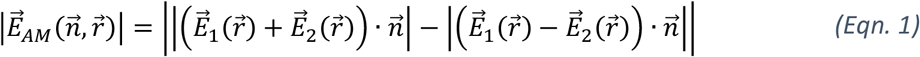

where *E*_1_and *E*_2_ are the E-fields generated by each electrode pair. At a given location, the maximum amplitude modulation along any direction is then obtained as:

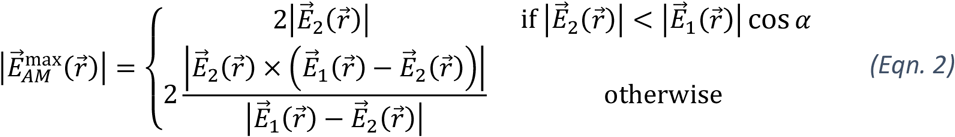

where *α* is the angle between the fields (assumed to be less than *π*/2), and the magnitude of *E*_1_ is, without loss of generality, assumed to be greater than the magnitude of *E*_2_^11^. The above metric is not supported by known mechanisms and is driven purely by phenomenological observations. Furthermore, despite a cosmetic similarity between the TIS exposure metric and the low-frequency field exposure magnitude, we caution the reader against such a comparison. The latter directly affects membrane polarization, while TIS must first undergo a demodulation process, the efficiency of which is unknown. We observe here that such mistaken comparisons occur frequently in the TI-related literature.

While the above metric is designed to characterize the stimulation-relevant low-frequency amplitude modulation exposure, a separate metric is needed for the high-frequency exposure. The frequency difference between the carrier signals leads to incoherent field superposition. The worst-case (highest) exposure occurs when the two fields are in phase and pointing in a similar direction:

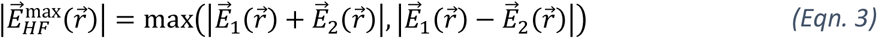

With regard to Joule heating, the incoherence means that the specific absorption rate (SAR) distributions from the two channels, rather than the E-fields themselves, must be summed to compute the heat source. That is, temperature increases are proportional to 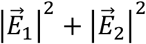 rather than 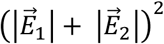, reducing the effect by up to a factor of two, if the two fields are identical except for their frequency.

### TIS and AEs

To date, there is little information regarding AEs specifically associated with TIS. Due to its recent emergence as a non-invasive modality for functional neuromodulation, the existing literature on TIS in humans is extremely sparse, with a preponderance of *in silico* modeling and simulation studies aimed at optimizing stimulation configurations^13,14,75,76^. However, Grossman et al.’s foundational 2017 study of TIS characterized its safety profile in rodents, including an immunohistochemical analysis of cellular and synaptic markers in the cortex and the hippocampus, and measurements of heating near the electrodes^11^. They found that relative to the unstimulated hemisphere and to a sham stimulation condition, TIS did not influence the number or density of apoptotic cells, nor did it negatively impact the physiology of microglia or astrocytes. In addition, synapse density was observed to be unaffected, and maximal heating did not exceed the largest spontaneous deviations from baseline^11^. Stimulation was conducted at 2 kHz and 2.01 kHz, 10 seconds on 10 seconds off, for 20 minutes with 125 *μ*A of current. Another animal study of TIS by Zhang et al., though less detailed in its evaluation of safety, provides further evidence that TI stimulation is safe in rodents^77^. The authors applied TIS in rats in 5 s blocks using a 2 kHz carrier signal, a modulation frequency of 3–10 Hz, and various total currents on the order of several mA. One week after the experiments, the modified neurological severity score (mNSS)^78^ was administered to assess neurobehavioral alterations and/or deficits. All rats were found to be normal in every aspect tested^77^.

In humans, a recent study by Piao et al. sought to characterize the safety profile of TIS for brain stimulation in a single-blind parallel (sham-controlled) trial with n=38 healthy volunteers^79^. Participants received a total of 30 min of active/sham stimulation in 10 min blocks, with the active arm (n=19) receiving either 20 Hz amplitude-modulated TIS (2 mA, 2 kHz and 2.02 kHz; n=9) or 70 Hz amplitude-modulated TIS (2 mA, 2 kHz and 2.07 kHz; n=10). Eye-closed, resting-state EEG recordings were obtained for 2 minutes before and after each stimulation block. Furthermore, serum neuron-specific enolase (NSE) was measured before and after stimulation, and electrode temperature was monitored during stimulation. Finally, the following neurological/neuropsychiatric tests were administered immediately before and after stimulation: the Montreal Cognitive Assessment (MoCA), the Purdue Pegboard Test (PPT), an abbreviated version of the California Computerized Assessment Package (A-CalCAP), a revised version of the Visual Analog Mood Scale (VAMS-R), a self-assessment scale (SAS), and a questionnaire regarding AEs. The authors found no significant differences between sham and active stimulation groups for the EEG band powers, serum NSE concentrations, and all neurological and neuropsychiatric evaluations. Furthermore, the relative change in all measurements were assessed *within* groups before and after stimulation, and found to be non-significant. Finally, the temperature at the skin-electrode interface remained well below body temperature for the duration of the experiment (range: 25.6–35.3 °C; median: 29.8 °C). Regarding AEs, several were reported though all were mild to moderate, and there were no significant differences between sham and active conditions. These included: mild itching, mild headache, mild warmth, mild tingling, fatigue, and vertigo^79^. Thus, the authors conclude that based on their findings, TIS is unlikely to induce neurological/neuropsychological state changes, or result in any adverse effects in humans for the range of stimulation parameters evaluated. This conclusion is informally supported by personal correspondence with collaborators, who report that TIS is well tolerated in humans, with no or minimal AEs similar to, but less frequent/intense than, tACS. The high frequencies employed in TIS mean that a primary limitation of tACS, namely, cutaneous sensation, is minimized. As the profile of TIS as a viable stimulation paradigm continues to increase, more human trials will be published, helping to clarify the landscape of TIS-specific AEs.

## Computational Characterization and Comparison of Electric Brain Stimulation Modalities

The mechanisms of electrical stimulation as well as potential associated health risks depend both on the stimulation settings (EM exposure, pulse shape, stimulation intensity) and on the electrophysiological, thermal, and electrochemical sensitivity/stimulability of the exposed brain tissues. Exposure distributions can vary considerably across stimulation modalities and electrode configurations. To help clarify this variation and facilitate an appraisal of TIS safety, we conducted a qualitative, comparative analysis of exposure metrics for tACS, tDCS, DBS, and TIS based on computational modeling in an anatomically detailed human head model (see *<u>Mechanistic Hypotheses and Relevant Dosimetric Quantity</u>* and *Appendix B* for details concerning metric selection). To this end, generic but representative setups for each modality were modeled numerically using EM and thermal simulations with realistic current strengths. On this basis, we inferred which of the application-specific safety concerns likely apply to TIS. Our investigation explored the effects of frequency and electrode shape and placement on heating and charge accumulation in brain and skin tissues.

All simulations were performed using the Sim4Life (ZMT Zurich MedTech AG, Switzerland) platform for computational life science, in conjunction with the MIDA^80^ reference anatomical human head model, developed by the IT’IS Foundation in collaboration with the FDA for computational investigations^81^. The MIDA model, generated from multimodal image data, includes dura and skull layers, and various stimulation targets. EM simulations were conducted using Sim4Life’s low-frequency electro-quasistatic and ohmic-current-dominated solvers (structured finite element method) with Dirichlet voltage boundary conditions at the active electrodes, and current normalization. Isotropic conductivity was assumed for all tissues. Conductivity magnitudes for each tissue were assigned using the IT’IS low-frequency database^82^. The computational domain (electrodes and head) was discretized using a rectilinear grid with an isotropic resolution of 0.5 mm in the brain and the tES electrodes, and 0.2 mm for the DBS electrode. E-fields were normalized to a 1 mA input current (per channel, in the case of TIS) to ensure meaningful field comparisons. Worst-case cathodic AF distributions were estimated using the largest absolute magnitude eigenvalue of the Hessian matrix of the electric potential^83^.

The following exposure setup conditions were selected for simulation:

- tES (tACS, tDCS and TIS require equivalent electro-quasistatic simulations): A typical setup comprising two circular electrodes, each with a surface area of 1.5, 3, 25 or 50 cm2, and an input current of 1 mA, was simulated as described above. E-field, current density, and AF all scale linearly with the total current, while power deposition and temperature increase scale quadratically. For sinusoidal currents, root-mean-square (RMS) and peak E-field and current density differ by a factor of 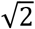 (RMS and peak quantities for non-sinusoidal exposures differ by other amounts). Here, we report peak values, reflecting typical clinical practice (however, note that reporting across literature is not consistent). Thus, the deposited power and concomitant temperature increase for tACS is twice that of tDCS.
- DBS: The E-field exposure in conventional DBS was calculated in the MIDA head model using a simplified model of the Medtronic 3389 DBS lead oriented along a realistic implantation trajectory with the two inner electrodes centered in the GP. Stimulation was achieved using a bipolar electric current applied between the two inner electrodes. Active electrode surfaces were assigned Dirichlet boundary conditions, with voltages calibrated to result in 1 mA of input current (passive electrodes were modeled as perfect conductors). Dielectric tissue properties were identical to those used in tES and TIS simulations.

### Peak E- and J-Field Strengths

Figure 2 depicts the E-field distributions produced by (top) a four contact, bipolar DBS electrode (modeled after Medtronic’s 3389 neurostimulator), and (bottom) a tES setup for three different diameters of circular electrode (fields normalized to 1 mA peak input current).

**Figure 2.**
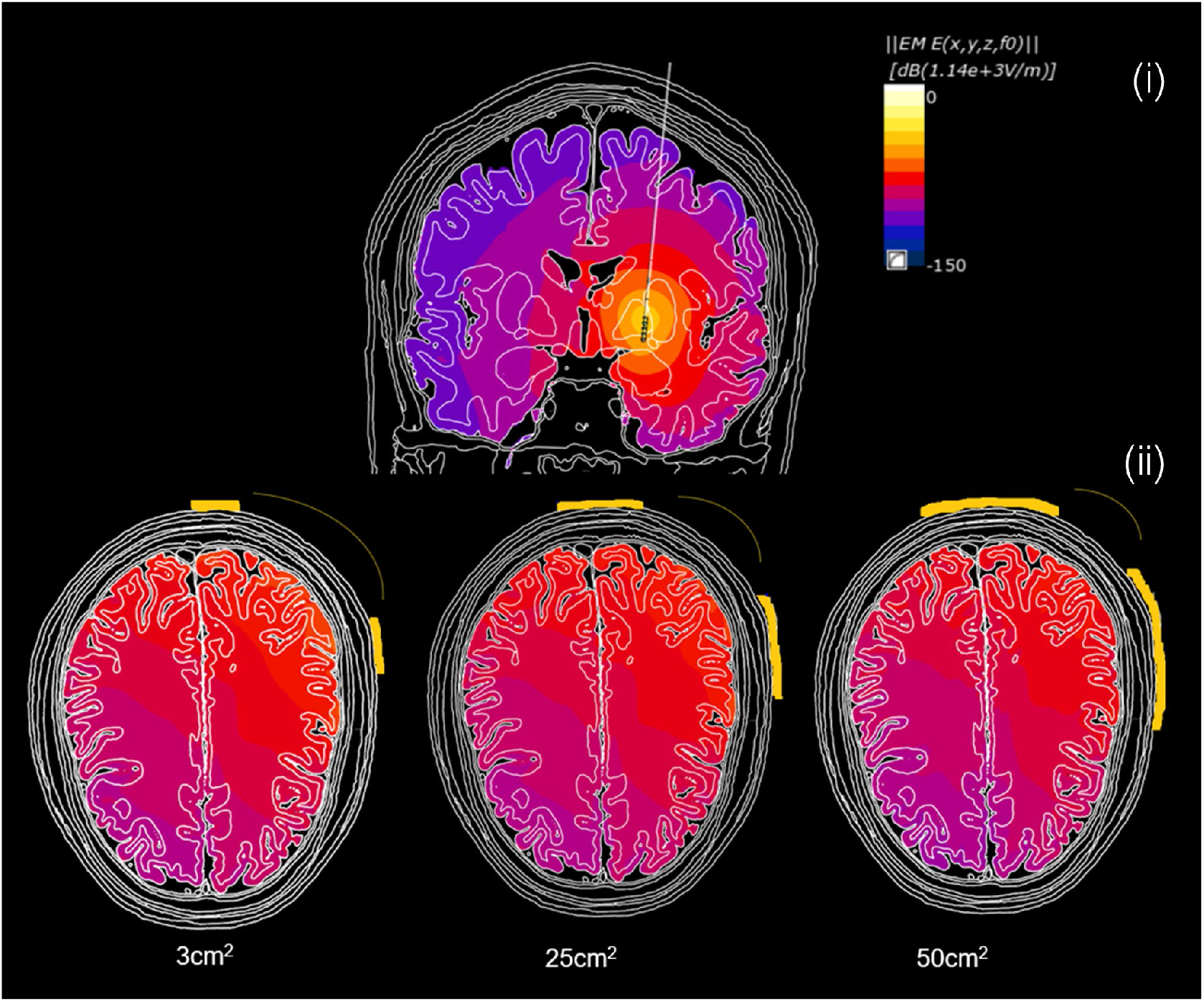
(i) Induced E-field distributions for a DBS electrode in bipolar configuration at 1 mA of input current (cf. Medtronic 3389 neurostimulator). (ii) tES field distributions for electrode placements identical to pair 1 in Figure 3, computed for circular electrodes with surface areas of 3, 25 and 50 cm^2^ and a 1 mA input current. Scale: 0 to -150 dB (reference: max value in DBS simulation; 10 dB per color step – one order of magnitude in field strengths for two steps) was chosen, demonstrating relatively minor brain exposure differences between different patch size when compared to DBS).

Spatial field peaks are reported separately for skin (in view of skin sensations) and brain (stimulation safety and efficacy). We applied 2 mm averaging to field values (E-field and current density (J-field)), as proposed by the ICNIRP 2010 guidelines, to obtain macroscopically relevant quantities and avoid numerical artifacts caused by the volume discretization or by segmentation-related sharp dielectric contrasts^84^. 2 mm averaging is suitable for external current sources, but poorly captures the much stronger field gradients near small, implanted electrodes. For point-like sources (spherically symmetric sources, *1/r*^*2*^ distance dependence), averaging has an impact below 20% at *r* >8 mm, while this occurs already at *r* >4 mm for line-like sources (cylindrical symmetry, *1/r* field dependence, e.g., near sharp electrode edges; note: for a DBS lead, that distance r does not necessarily correspond to the distance to the lead center line, but can be the distance to a sharp electrode feature, such that the divergence can be situated in close contact to the tissue). At a distance of 2 mm (0.8 mm for line-like sources), 2 mm averaging can already cause two-fold differences over baseline; in our simulations, the unaveraged peak values at the DBS electrode are as high as 1110 V/m/mA due to strong field gradients, while unaveraged peak tES exposure in the brain is 2.3 V/m/mA and 80 V/m/mA in the skin. Hence, relying on averaged quantities in close proximity to the electrode can be misleading.

For this analysis, we define the region of “activated” brain tissue to be the volume in which the 2 mm averaged E-field is >90% of its peak value. The activated brain volume was 1.4 mm^3^ for tES (3 cm^2^ electrode size; peak: 1.6 V/m), and 7 mm^3^ for DBS (peak: 145 V/m). Thus, peak intracranial E-fields induced by DBS were about two orders of magnitude greater than those produced by tES, as were the field gradients, with a factor of 40 difference in focality. With regard to current density, for DBS we found a 2 mm line averaged J-field peak of 35 A/m^2^ localized in the GP near the electrodes. For tES exposure, we found a 2 mm line averaged J-field peak of 0.4 A/m^2^ in the brain. In the skin, the corresponding quantities for DBS and tES were 0.002 A/m^2^ and 5.1 A/m^2^, respectively.

### TIS

The location and magnitude of the maximum E-field amplitude modulation provide information about the likelihood of TIS neuromodulation, while the combined E-field strength governs safety-relevant physical interactions with the tissues (e.g., thermal heating). To ensure safe and effective TIS, the spatial distribution of both the TI amplitude modulation and the total E-field strength must be estimated to minimize off-target stimulation and AEs. Figure 4 compares the distributions of total and directional maximum amplitude modulation (amplitude modulation of the projection along the local principal fiber orientation obtained from DTI data), as well as the total and directional combined high-frequency E-fields for a generic TIS setup. Note that the TIS electrode montage and steering parameters were not optimized for a specific target, and that no effort was made to avoid hotspots. The current ratio was fixed at 1:3 (blue electrode pair: 0.5 mA; red electrode pair: 1.5 mA).

**Figure 3.**
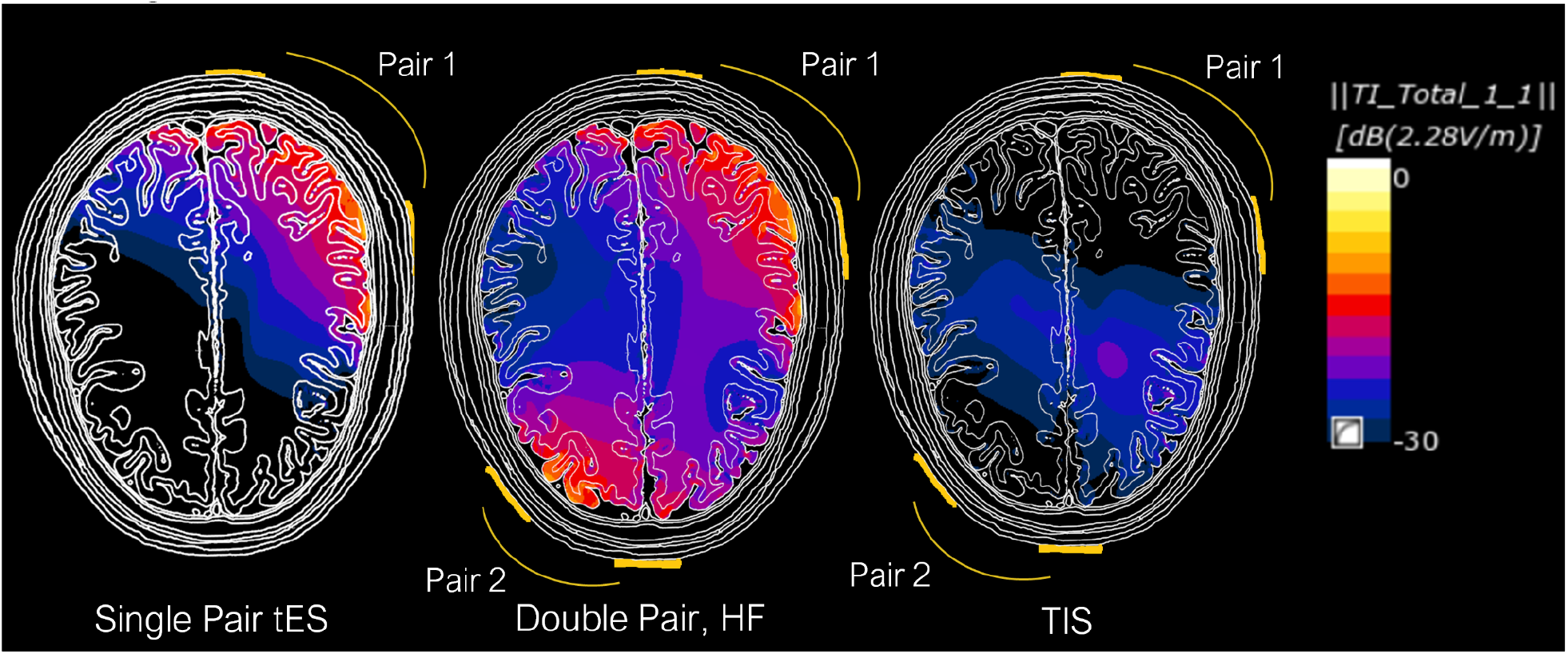
Comparison between conventional single pair tES (left) and total TIS high frequency E-field exposure (middle), as well as the corresponding low-frequency TIS modulation magnitude distribution (right). The total TIS carrier frequency E-field map (middle) shows the maximal high frequency field magnitude achieved for in-phase, constructive interference.

**Figure 4.**
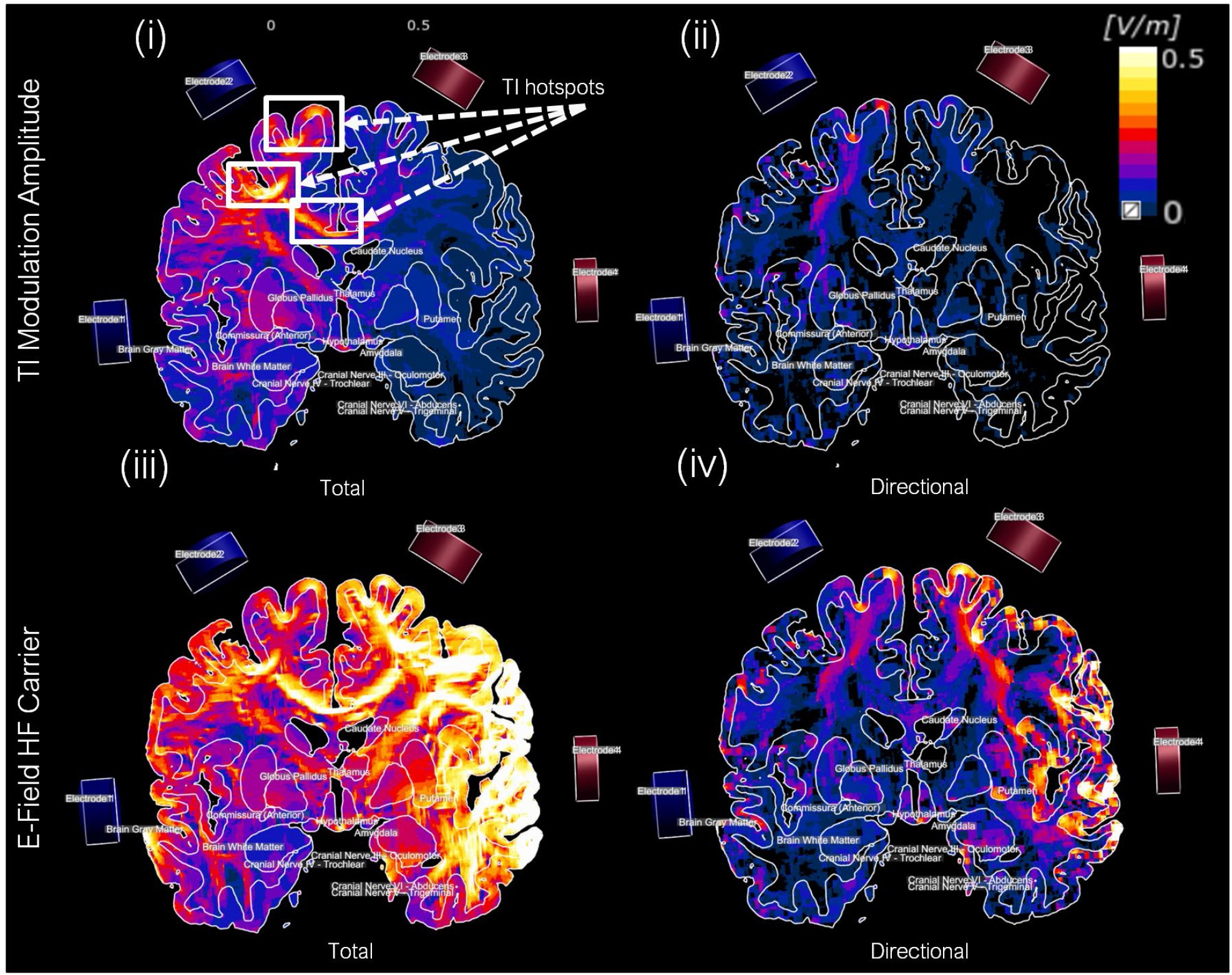
TI amplitude modulation magnitude and combined high-frequency E-field distributions (sagittal slice views). Note that the electrode montage and steering parameters have <u>not</u> been optimized for a specific target and that no effort has been made to reduce exposure hot spots elsewhere. Blue electrodes, 0.5 mA; red electrodes, 1.5 mA (current ratio of 1:3). (i) Maximum (along any orientation) TI amplitude modulation magnitude. (ii) Directional TI amplitude modulation magnitude projected along the DTI-derived principal local fiber orientation. (iii) Combined high-frequency E-field (i.e., total field magnitude for in-phase channel contributions). (iv) Directional projection of the combined high-frequency E-field (i.e., high-frequency field component projected along the principle orientation of local neural tracts).

As the low-frequency modulation amplitude is dominated by the weaker field, it is maximal in regions where the two channels’ exposures are both high and of similar magnitude, permitting deeper and more localized targeting. However, TI remains subject to physical constraints – i.e., TI exposure peaks (foci) are typically subsets of the high-frequency peaks and it can be difficult to avoid high TI exposure at locations with dielectric features such as nearby CSF (ventricles, sulci) that bundles a guides current flow, conductive implants, or conductivity heterogeneity (e.g., dielectric contrast) and anisotropy. Tissue structure (e.g., strongly preferential fiber/neuron orientations) may further increase TI exposure localization, which can help or hinder in targeting the desired structure. To avoid TIS of overlying structures, either the electrodes must be sufficiently separated, or E-fields in the off-target structures must be nearly perpendicular. See Figure 4 for a depiction of the shift in TI modulation amplitude towards the electrode pair with the weaker current (blue pair).

### Thermal Increase

Thermal simulations were conducted in the MIDA head model based on the Pennes Bioheat Equation (PBE)^85^:

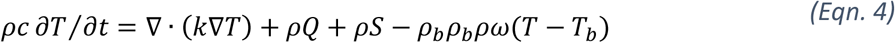

where *ρ* and *c* are the density and the specific heat capacity distributions, *k* is the thermal conductivity, *Q* the specific metabolic heat generation rate, *S* the SAR (obtained from coupled EM simulations), *ω* the perfusion rate, and *ρ*_*b*_, *c*_*b*_ and *T*_*b*_ are the density, specific heat capacity, and temperature of the arterial blood. Instead of solving for the absolute tissue temperature, we used the PBE to calculate the temperature increase *ΔT* with respect to the steady-state temperature distribution. This is possible so long as the temperature dependence of perfusion can be ignored. In addition to improving numerical accuracy, this approach avoids the need to consider metabolic heat generation, external temperature, and arterial blood temperature. Simulations were performed with Sim4Life’s stationary PBE solver using the time-averaged power density as a heat source (DBS pulsation is much faster than the thermal time-constant). Tissue thermal properties were assigned from the IT’IS database of thermal and dielectric tissue properties^82^. Convective boundary conditions were applied at the interface with the external air and internal airways (convection coefficient *h=*30 W/m^2^/K internally, *h=*6 W/m^2^/K on the external surface) to account for active convection. In the presence of multiple currents (e.g., TIS channels), coherent field superposition was used for identical frequencies, and incoherent superposition (i.e., SAR addition) was used when the frequencies differed.

Figure 5 illustrates the steady-state temperature increase *ΔT* distribution for 1 mA of input current. A peak temperature increase of *ΔT*/*I*^2^=0.026°C/mA^2^ was found for the DBS implant, while an increase of 0.002°C/mA^2^ was found for the tDCS setup using the smallest electrode size (3 cm^2^). For larger electrodes, the temperature increase was smaller than 0.001°C/mA^2^ or smaller than 0.1°C for 10 mA. The temperature increased roughly in inverse proportion to the electrode area. In both cases, heating was confined to an area near the electrodes, and was well below the threshold for direct tissue damage.

**Figure 5.**
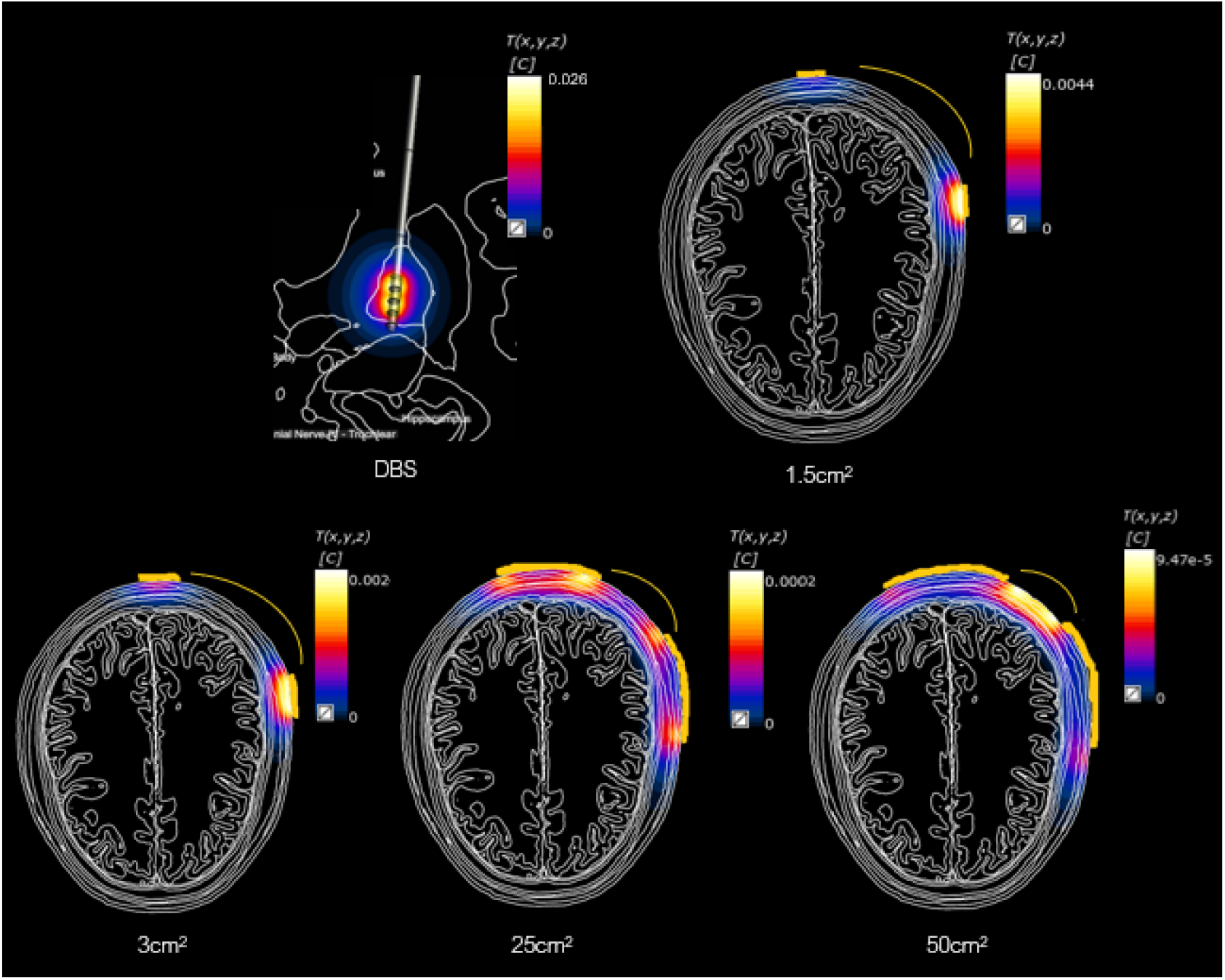
Simulated steady-state temperature increase (*ΔT*) distributions for DBS using a bipolar electrode configuration (top left), and tES for an input current of 1 mA and varying electrode sizes. Heating is principally localized near the electrodes, such that brain heating is minimal for tES. In all cases, the heating is well below thresholds for direct tissue damage.

### AF and Volume of Activated Region

AF peak magnitude data must be interpreted with care, since: (i) the AF distribution can show divergent behavior near sharp electrode features, which also results in strong dependence on the discretization resolution and shape of the electrode, (ii) the AF is not well defined at dielectric interfaces (such as the grey-matter/white matter interface that is believed to be a highly relevant interaction site); and (iii) even within a single tissue, such as white-matter, heterogeneity and anisotropy strongly affect the AF and are difficult to capture in computational models. In brain stimulation applications, the AF is primarily valuable for mechanistic investigations, and the prediction of the volume of tissue activated (important for DBS device design)^86^. With the above-mentioned limitations in mind, Figure 6 shows qualitative AF distributions in the brain for DBS and tACS, obtained by spectral decomposition of the Hessian matrix of the electric potential, from which we extracted and plotted the maximal absolute eigenvalues (corresponding to the most stimulable orientation in biphasic exposures)^†^. Also, contributions to the AF due to dielectric contrast at the interfaces between tissues were omitted in this calculation, as was the influence of brain tissue heterogeneity, meaning that these data provide only a qualitative picture of stimulation likelihood. The actual AF threshold for neurostimulation depends on fiber type and pulse shape. As the figure illustrates, AF-related neurostimulation effects are significantly stronger for DBS than for tES.

**Figure 6.**
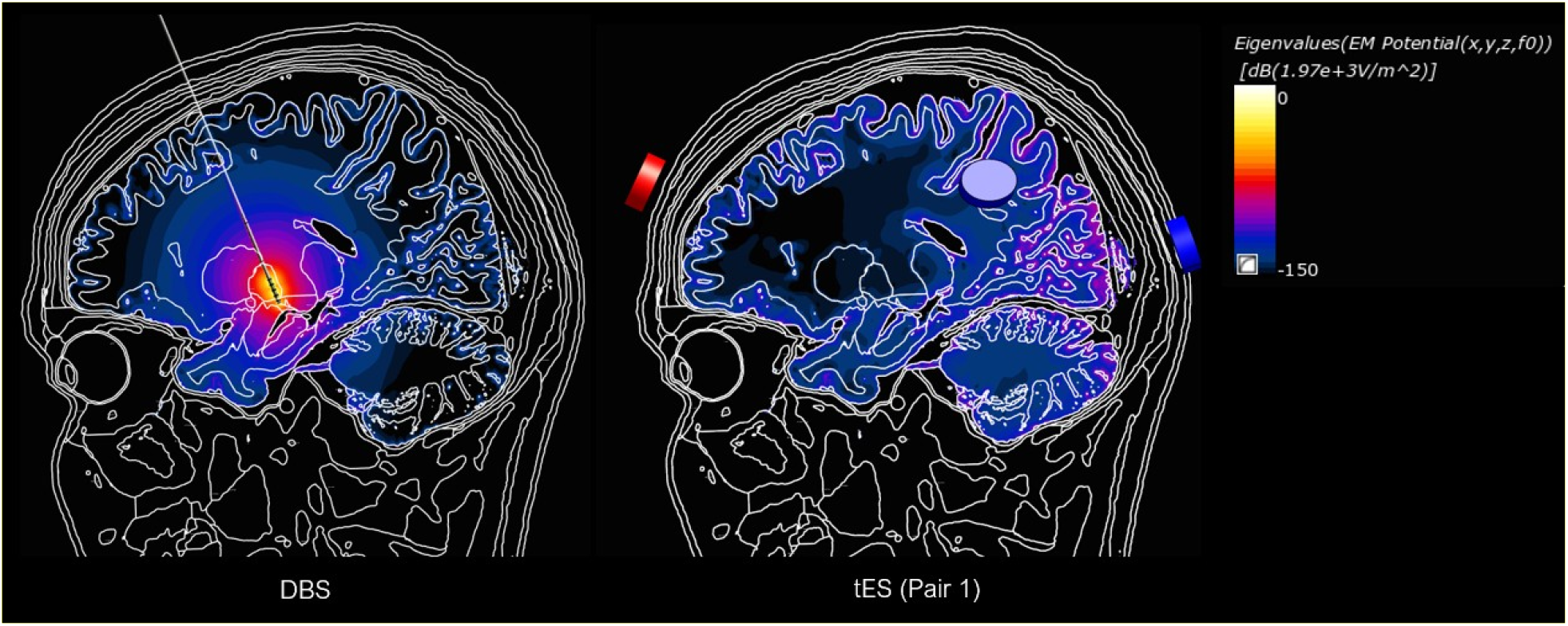
Cross-section images of the maximal Hessian eigenvalue distributions for the electric potential, a measure of the maximal normalized AF of arbitrarily oriented straight fibers (left: bipolar DBS; right: tES; current magnitude: 1 mA; electrode size: 3 cm^2^).

## Comparison of Exposure Metrics

The results of the biophysical exposure simulations are summarized in Table 2, which constitutes the basis for comparing TIS with established neurostimulation modalities to propose safety thresholds:

**Table 1.** Overview of common DBS stimulation parameters (for more details see Table A.3 in *Appendix A*).

| Quantity | Min | Max |
| --- | --- | --- |
| repetition frequency | 2 Hz | 333 Hz |
| voltage | 0.4 V | 14 V |
| current | 0.005 mA | 12 mA |
| pulse duration | 15 μs | 520 μs |

**Table 2.**
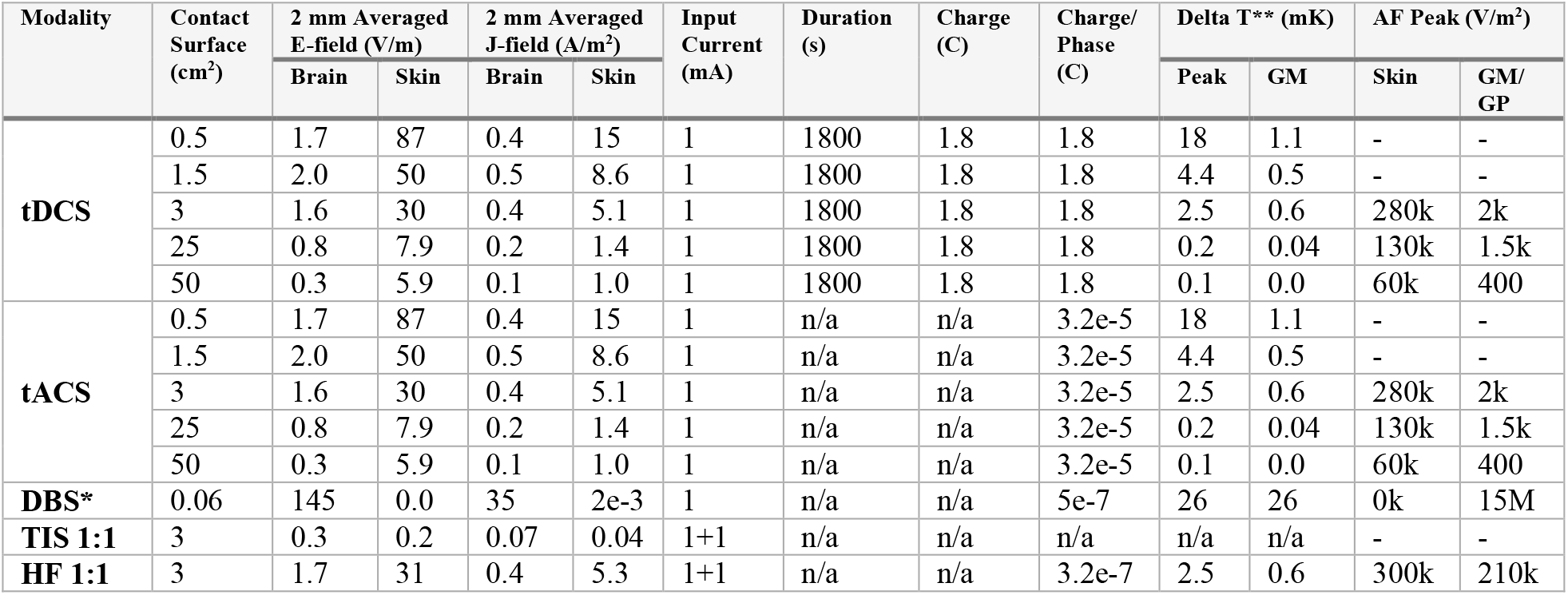
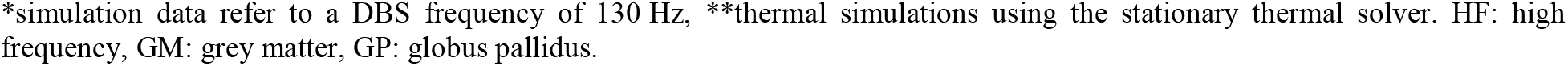
Dosimetric and thermal exposure quantities for typical tDCS, tACS, DBS, and TIS configurations. All exposures assume an input current of 1 mA. For tACS and DBS, the charge is calculated per phase (e.g., half-period for sinusoidal currents).

| Modality | Contact Surface (cm<sup>2</sup>) | 2 mm Averaged E-field (V/m) |  | 2 mm Averaged J-field (A/m<sup>2</sup>) |  | Input Current (mA) | Duration (s) | Charge (C) | Charge/Phase (C) | Delta T** (mK) |  | AF Peak (V/m<sup>2</sup>) |  |
| --- | --- | --- | --- | --- | --- | --- | --- | --- | --- | --- | --- | --- | --- |
|  |  | Brain | Skin | Brain | Skin |  |  |  |  | Peak | GM | Skin | GM/GP |
| tDCS | 0.5 | 1.7 | 87 | 0.4 | 15 | 1 | 1800 | 1.8 | 1.8 | 18 | 1.1 | - | - |
|  | 1.5 | 2.0 | 50 | 0.5 | 8.6 | 1 | 1800 | 1.8 | 1.8 | 4.4 | 0.5 | - | - |
|  | 3 | 1.6 | 30 | 0.4 | 5.1 | 1 | 1800 | 1.8 | 1.8 | 2.5 | 0.6 | 280k | 2k |
|  | 25 | 0.8 | 7.9 | 0.2 | 1.4 | 1 | 1800 | 1.8 | 1.8 | 0.2 | 0.04 | 130k | 1.5k |
|  | 50 | 0.3 | 5.9 | 0.1 | 1.0 | 1 | 1800 | 1.8 | 1.8 | 0.1 | 0.0 | 60k | 400 |
| tACS | 0.5 | 1.7 | 87 | 0.4 | 15 | 1 | n/a | n/a | 3.2e-5 | 18 | 1.1 | - | - |
|  | 1.5 | 2.0 | 50 | 0.5 | 8.6 | 1 | n/a | n/a | 3.2e-5 | 4.4 | 0.5 | - | - |
|  | 3 | 1.6 | 30 | 0.4 | 5.1 | 1 | n/a | n/a | 3.2e-5 | 2.5 | 0.6 | 280k | 2k |
|  | 25 | 0.8 | 7.9 | 0.2 | 1.4 | 1 | n/a | n/a | 3.2e-5 | 0.2 | 0.04 | 130k | 1.5k |
|  | 50 | 0.3 | 5.9 | 0.1 | 1.0 | 1 | n/a | n/a | 3.2e-5 | 0.1 | 0.0 | 60k | 400 |
| DBS* | 0.06 | 145 | 0.0 | 35 | 2e-3 | 1 | n/a | n/a | 5e-7 | 26 | 26 | 0k | 15M |
| TIS 1:1 | 3 | 0.3 | 0.2 | 0.07 | 0.04 | 1+1 | n/a | n/a | n/a | n/a | n/a | - | - |
| HF 1:1 | 3 | 1.7 | 31 | 0.4 | 5.3 | 1+1 | n/a | n/a | 3.2e-7 | 2.5 | 0.6 | 300k | 210k |

\*simulation data refer to a DBS frequency of 130 Hz, \*\*thermal simulations using the stationary thermal solver. HF: high frequency, GM: grey matter, GP: globus pallidus.

For the same electrode montage and total current, the combined field exposure magnitude (carrier) for TIS is less than or equal to that of tACS, with more spatial dispersion, as a consequence of being distributed across a greater number of electrodes. However, TIS carrier frequencies are one to two orders of magnitude higher, making it far less likely to result in direct neuromodulation. Both tACS and TIS (at least when using only two high-frequency carriers) have diminished peak field strengths and reduced focality in the brain in comparison to DBS. The low-frequency modulation envelope magnitude distribution of TIS is more localized than that of tACS, but as it is dominated by the weaker field of the two channels, it is never larger than the tACS field, and its local maxima/foci are typically a subset of those generated by tACS. We also note that the stimulation efficiency of the TI modulation magnitude should not be compared directly to tACS field strengths at the same frequency, since TIS requires an additional demodulation step and the mechanisms of action are different. All modalities investigated resulted in minimal heating that would not be expected to cause direct thermal tissue damage (see *Appendix B*). With respect to heating, charge, current and interface effects, TIS is comparable to tACS.

In summary, the procedural aspects of TIS and tACS are largely overlapping, despite certain differences; TIS is more focal, uses high frequency carriers that are less likely to elicit direct neurostimulation near the electrodes, and is capable of modulating deeper brain structures. Compared to DBS, which also targets deep brain regions with enhanced focality, TIS is nearer to tACS in terms of the magnitude and size of local maxima. Thus, safety considerations for tACS also broadly apply to TIS, with two additional TIS-specific concerns: (i) in TIS, the higher frequencies of applied currents increase the magnitude threshold required to evoke skin sensations, which would otherwise serve as cautionary signals, and (ii) TIS may entail the use of electrode montages that would normally be avoided owing to the possibility of activating nearby cortical or cerebellar structures. Furthermore, the biophysical mechanisms of TIS are still poorly understood, warranting a degree of general caution.

### Additional Considerations for Frequencies >10 kHz

The possibility of using very high frequency carriers (on the order of 10–100 kHz) for TIS merits discussion, though the capacity of neurons to demodulate such fields is as of yet unknown. In this frequency range, direct effects resulting from exposure to high frequency fields are expected to be further diminished relative to lower frequencies, as safety guidelines specify current thresholds proportional to frequency^84,87^. However, at such high frequencies, the relative importance of capacitive currents (compared to ohmic currents is) may no longer be negligible, and additional simulations accounting for the frequency dependence of dielectric materials would be needed to quantify whether and how the exposure quantities summarized in Table 2 are affected.

### Electrode Size, Shape, and Placement

To investigate the impact of electrode size and separation distance on tissue exposure under highly controlled conditions, we developed a simplified, cylindrical head model (see Figure 7) comprising the following tissues: skin, SAT, galea, skull (three layers: cortical inner and outer table with cancellous diploe in between), dura, CSF, gray and white matter. Table 3 summarizes the tissue dielectric and geometric properties. Circular electrodes with varying surface areas (1.5, 3, and 25 cm^2^) were placed on the head model as shown in Figures 7 and 8. This arrangement was discretized using a homogeneous rectilinear grid with 0.2 mm resolution. Field distributions were calculated for varying inter-electrode separations, employing Sim4Life’s ohmic current-dominated low-frequency EM solver with Dirichlet voltage boundary conditions at the active electrodes (inactive electrodes were excluded from the simulations), and with total current normalized to 1 mA. Peak values for current density and E-field magnitude were reported for each simulation condition over the set of all combinations of electrode sizes and separations.

**Table 3.** Head model parameters: thicknesses and electric conductivities^82^.

| Tissue Name | Thickness [mm] | Conductivity [S/m] |
| --- | --- | --- |
| Skin (Epidermis and Dermis) | (0.07+1.8) | 0.17 |
| SAT | 3 | 0.057 |
| Galea | 0.8 | 0.368 |
| Cortical Bone (Outer) | 2.6 | 0.0035 |
| Cancellous Bone | 2.0 | 0.082 |
| Cortical Bone (inner) | 2.6 | 0.0035 |
| Dura | 0.6 | 0.461 |
| CSF | 1 | 1.71 |
| Gray Matter | 2.5 | 0.24 |
| White Matter | 15.4 | 0.265 |

**Figure 7.**
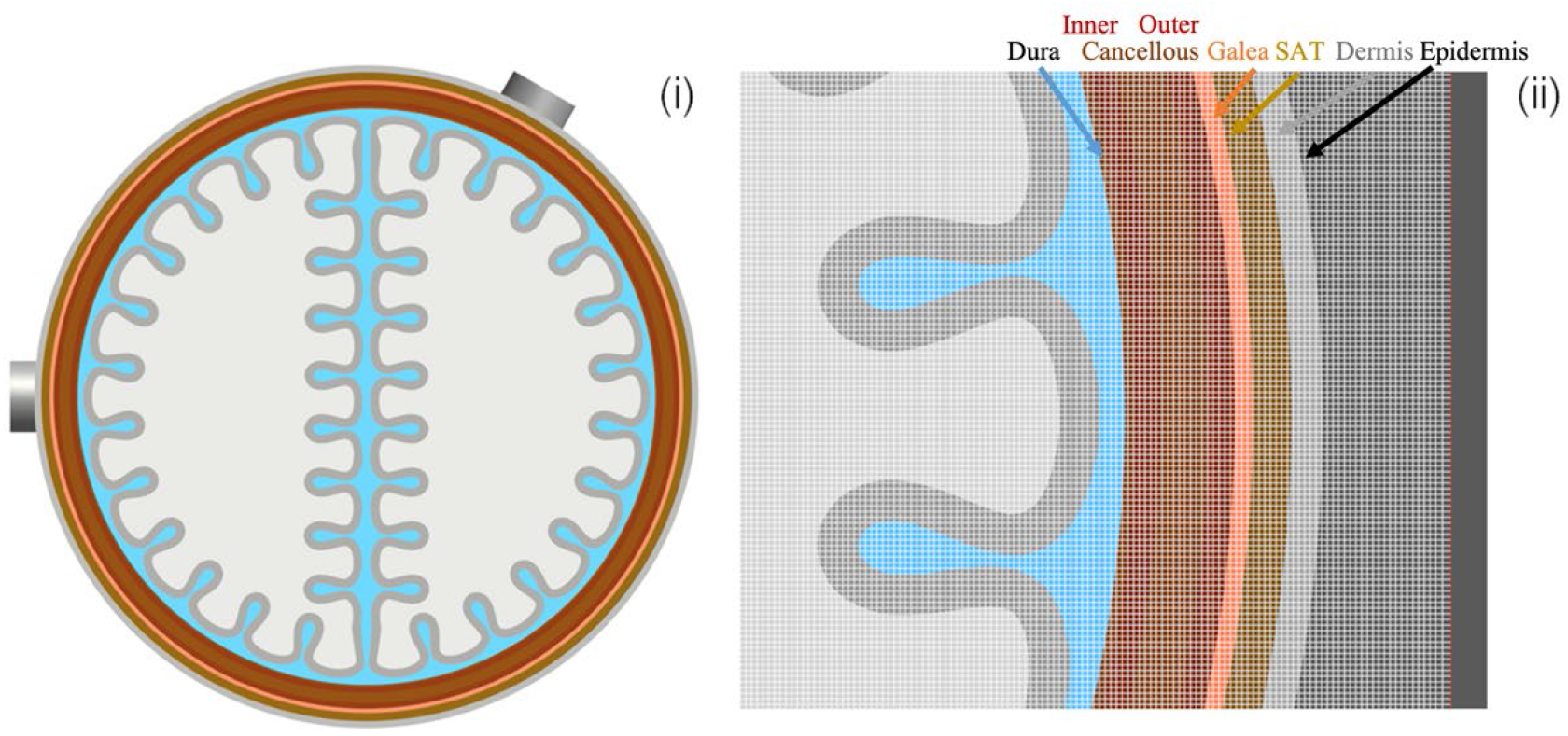
(i) Geometry of the simplified, cylindrical head model, with detail of the tissue layers: epidermis, dermis, SAT, galea, three skull layers (cortical inner and outer table, cancellous bone), dura, CSF, gray and white matter. The external diameter of the head model is 18.6 cm and the external diameter of the white matter is 15.4 cm. The height of the cylinder is 12 cm. Electrodes for every condition were placed as illustrated in (i). (ii) Detail of the head model, along with the rectilinear grid used for discretization.

**Figure 8.**
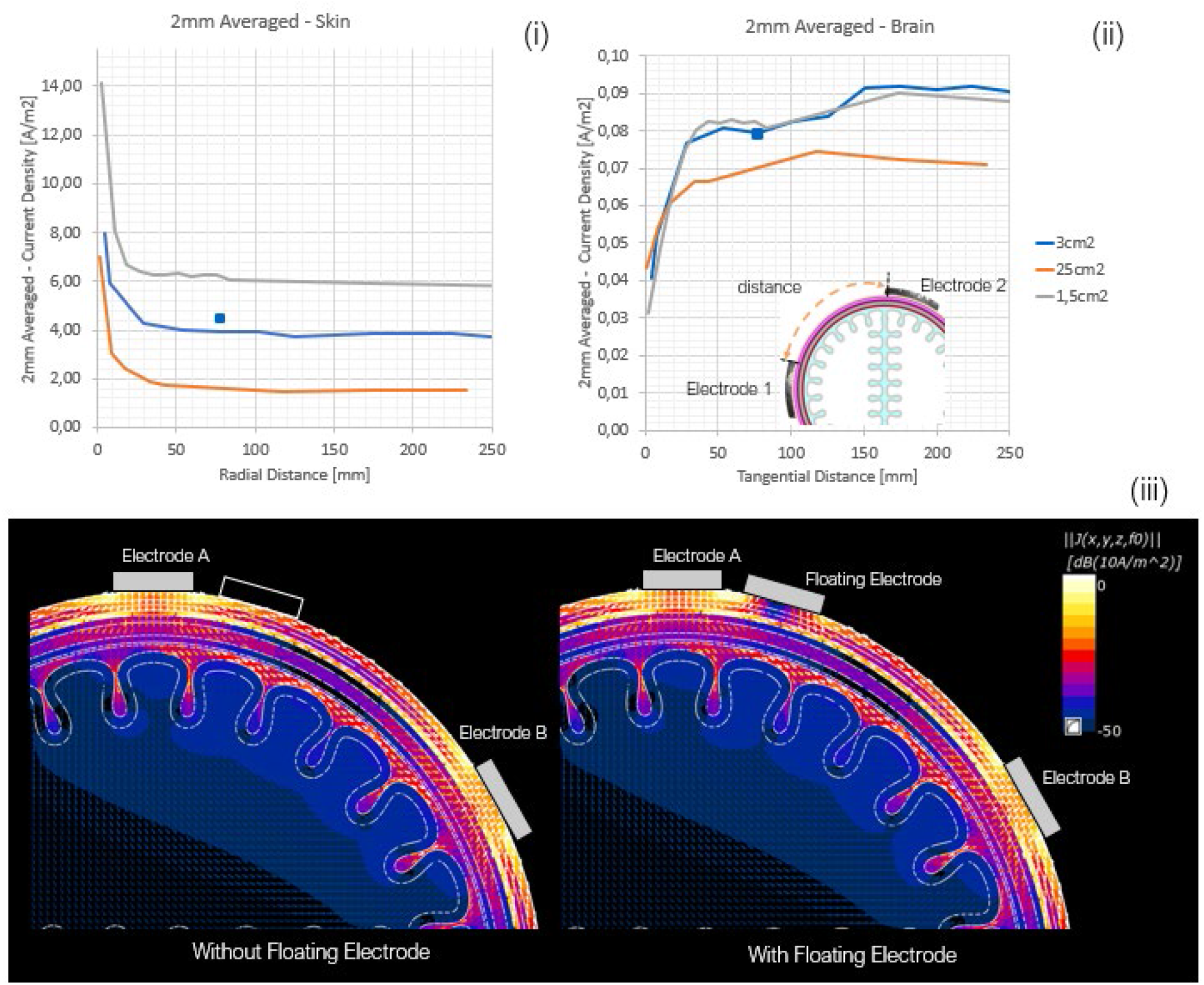
2 mm averaged current density in (i) skin and (ii) brain (gray matter) of the simplified head model for three different electrode sizes (1.5, 3, and 25 cm^2^) as a function of the inter-electrode separation (length of the shortest geodesic path on the skin surface). Small blue squares: 2 mm averaged peak current density for simulations with a floating electrode that does not contribute current (shown in (iii)). (iii) Current density distribution for 3 cm^2^ electrodes in the absence (left) and presence (right) of an additional electrode that does not provide current but may enhance the field as a result of edge effects.

The results indicate that a large fraction of current flows through the scalp (primarily through muscle and galea), while the portion that enters the brain must cross twice through the skull with its highly resistive bone layers. The amount of current reaching the brain is further attenuated by shunting of the skull-crossing current through the CSF. The magnitude of this effect is a function of CSF continuity, which may change depending on the subject’s posture. Cortical gyrification also affects the path of current flow and E-field orientation. E-field enhancement is apparent at the electrode border, which is a result of the sharp conductor edge and strong local curvature of field lines. In addition, we found that the influence of electrode separation on peak E-field and current density falls below 20% above a distance of approximately 50 mm. Peak brain exposure is dominated by skull resistance, and peak epidermis exposure by electrode edge effects. For the electrodes in close proximity, the edge effects of the two electrodes begin to merge, pushing local skin exposures higher, and driving decreases in brain exposure due to current shunting through the scalp. Electrode size is an important determinant of skin exposure since: (i) current density below the electrode is inversely proportional to the electrode area, whereas the importance of electrode size is diminished in the brain due to current dispersal, (ii) edge effects, which are safety and sensation limiting in the skin, are directly related to electrode edge length (edge field strength and current density are inversely proportional to edge length), and (iii) power deposition and heating, which are dominated by edge effects, scale with the square of field magnitude, and therefore in roughly inverse proportion to electrode area. The impact of electrode size on brain exposure, however, is less prominent. In summary, exposure strength is dependent on both external system parameters and intrinsic anatomy. The former is dominated by skin contact area (see *<u>Discussion</u>*) and electrode edge sharpness (which enhances local fields), while the latter is influenced chiefly by local skull resistance (which varies considerably between subjects), the continuity of CSF coverage, and local cortical folding.

Figure 9 shows the distributions and peak values of current density and E-field in the skin and brain for a fixed input current of 1 mA as a function of electrode separation (3 cm^2^ electrode area). The presence of passive electrodes, or electrodes operated at different frequencies (as required for TIS), can lead to important edge effects due to field concentration at sharp conductor features. Figure 8 illustrates this principle; E-field and current density in the scalp are markedly increased in close proximity to the electrodes in the presence of an additional electrode that does not provide current. In TIS, which requires the use of multiple electrodes operating at different frequencies, such effects may have safety implications, or cause skin sensations that affect blinding or trigger functional and/or behavioral responses.

**Figure 9.**
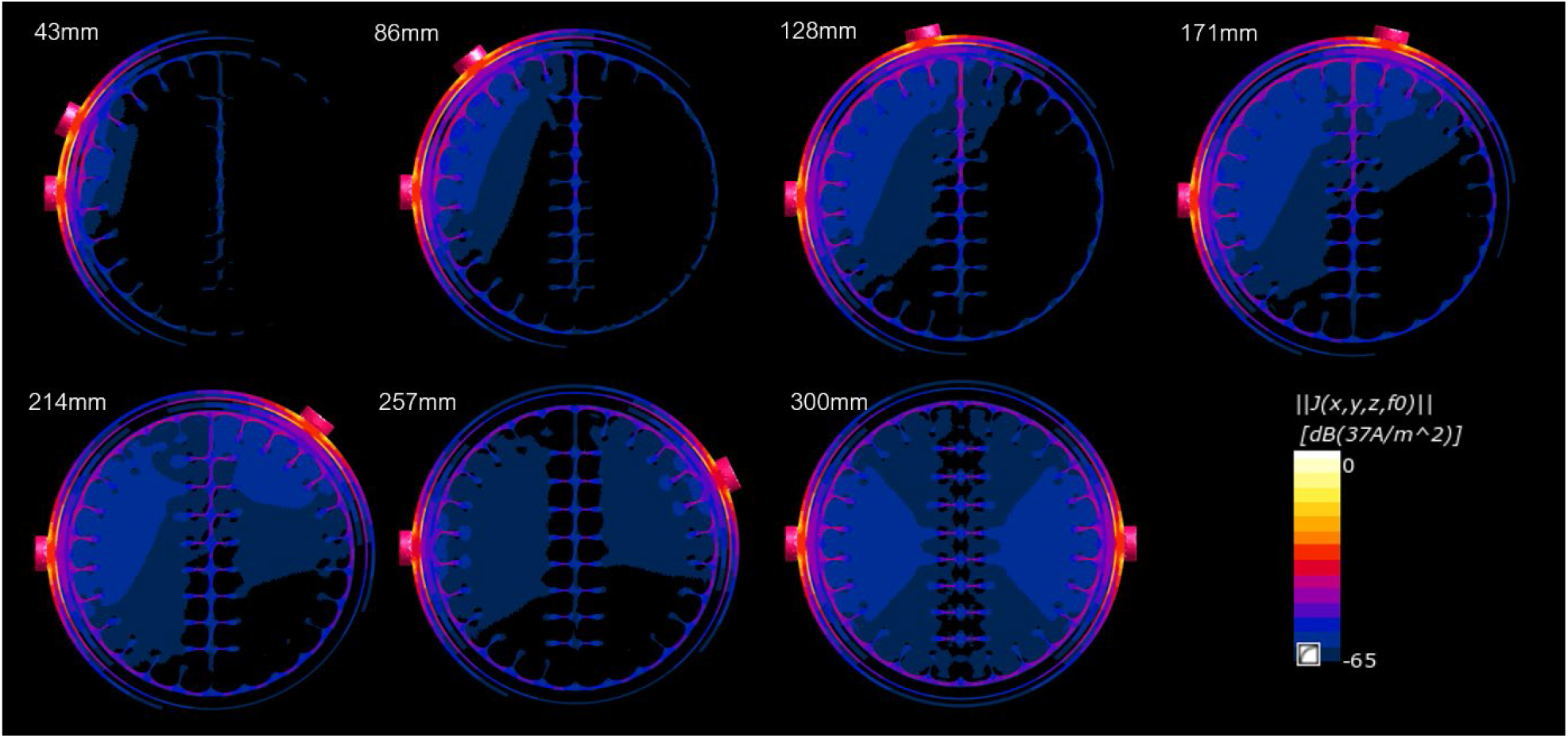
Current density distributions for increasing electrode separations. Simulations were conducted using a simplified head model, 3 cm^2^ electrodes, and 1 mA of input current.

Local field enhancement and concentration is also an issue near invasively placed electrodes (either sensing or stimulating) and can result in unwanted stimulation or affect targeting. Consequently, no transcranial stimulation should be performed for subjects with highly conductive brain implants in direct proximity to neural tissue without an extensive and careful risk assessment.

### Threshold Proposal

On the basis of the analysis of computed exposure and interaction physics safety thresholds for TIS are proposed. These thresholds are intended to ensure safety, but do not guarantee sensation-free stimulation, which can disrupt study blinding, but does not pose a danger to health.

Modern tES stimulation protocols regularly utilize total currents on the order of 2 mA (see Table A.3, *Appendix A*, average current amplitude: 1.7 mA; range: 0.75–2 mA), and early work with TIS in human subjects has applied up to 4 mA of total current (up to 3 mA in a single channel), in the low kHz range without eliciting AEs (personal correspondence with collaborators). Assuming an electrode size of 1.5 cm^2^, this translates to peak 2 mm averaged E-field values in the brain of approximately 8 V/m in the low frequency range, and 5 V/m for kHz exposures. In the skin, the corresponding values are 80–200 V/m in the low frequency range and up to 90 V/m for kHz exposures. To better understand the effects of localized deep brain stimulation, we turn our attention to DBS. DBS targets (GP and STN) are well contained within a sphere of radius 15 mm (based on Medtronic 3389 DBS neurostimulator placed in the GP; sphere centered on the geometrical center of the four contacts), outside of which it may be assumed that DBS exposure does not directly affect brain activity. Typically, DBS employs voltages in the range of 5–14 V (see Table A.3, *Appendix A*, average voltage: 6.6 V), which produce currents of up to 20 mA (clinically reported impedances are in the 0.5–1.5 kΩ range^88^ while simulations typically report 0.6–1.2 kΩ^89^). The associated peak-averaged E-fields in the brain outside the sphere are roughly 7 V/m but can reach 30 V/m (see *Appendix B*).

We can gain a purchase on reasonable current thresholds for TIS by comparing TI-induced brain E-field magnitudes, the difference frequency modulation amplitude, and the total current per channel with standard exposure conditions for tES, and with peak off-target field magnitudes for DBS. This approach suggests that TIS exposures are likely safe for currents in the range of 3–5 mA at typical tES frequencies, and for currents spanning 15–30 mA for typical DBS frequencies of up to 200 Hz. Our analysis is limited by the sparse availability of data for low kHz tES, and the consequent lack of safe use history at higher exposure levels. Given the tradeoff between the strength and duration of threshold neurostimulation^90^, and in accordance with ICNIRP guidelines^†^, we propose the following structure for safe exposure limits: constant stimulation thresholds up to a chosen frequency (ICNIRP guidelines suggest 2.5 kHz; simulations of myelinated and unmyelinated A- and C-fibers suggest < 1 kHz), beyond which thresholds increase linearly with frequency. On this basis, TI currents up to 16 mA below 2.5 kHz, 30 mA at 4 kHz, and >500 mA at 100 kHz are theoretically acceptable. However, since tissue heating is proportional to the square of current, thermal safety considerations become relevant for the increased current thresholds allowed at high frequencies. For typical tES and DBS current magnitudes (and accounting for DBS interpulse intervals), skin and brain temperature increases remain in the mK range. Even the highest clinical DBS magnitudes only increase temperature nearby the implant by tenths of a Kelvin^91^. Our simulations show that current magnitudes below 15 mA result in peak temperature increases in the brain of no more than 0.2 K, which is considered to be safe. At these current strengths, skin heating is well below the 2 K threshold accepted by the FDA for medical applications. Even for the comparatively conservative ICNIRP^84^ and IEEE^87^ safety guidelines, currents on the order of 100 mA would be required to exceed the 1 K limit. Charge accumulation is not a limiting factor for TIS since the charge injected per phase is proportionally reduced with increasing frequency.

The safety thresholds described above are summarized in Table 4. These thresholds are intended to ensure safety, but do not guarantee sensation-free stimulation, which can disrupt study blinding but does not pose a danger to health. All quantities were computed for 3 cm^2^ TIS electrodes, reflecting common practice. Larger electrode areas would tend to increase current thresholds, while poor electrical contact (resulting in reduced contact area) or closely neighboring electrodes would reduce the thresholds for safe current exposure. Smaller electrodes may be used provided that the maximum current is reduced in proportion to the electrode contact area.

**Table 4.**
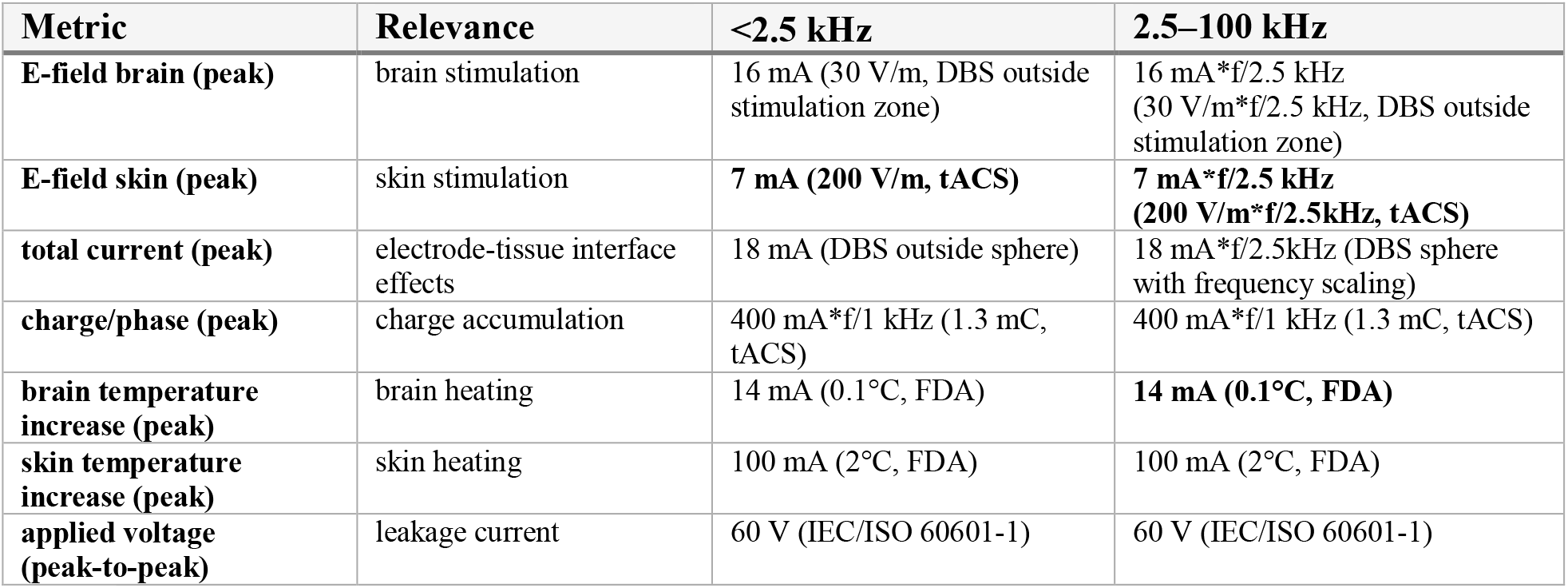
Proposed safety thresholds for TIS by exposure metric (3 cm^2^ electrodes). Selection of exposure mechanisms was motivated by reported AEs. Thresholds are based on mechanistic considerations, dosimetric simulations of tDCS, tACS, DBS and TIS, and the literature-informed history of safely applied conventional electrical stimulation. Thresholds are formulated in terms of measurable TIS application parameters (applied currents and voltages). Exposure quantities are shown in parentheses together with the reference application/standard used to define the threshold. All quantities are expressed as peak values (not RMS) except for applied voltage, which is formulated as peak-to-peak to ensure consistency with leakage current. Values in bold highlight the lowest effect thresholds and the metrics that may limit TIS exposure. All quantities were computed based solely on the direct effects of applied currents and voltages. IEC/ISO: joint technical committee of the International Organization for Standardization and the International Electrotechnical Commission. Stimulation zone: A sphere of radius 15 mm, centered on the DBS contacts (based on Medtronic 3389 neurostimulator) and encompassing the GP and the STN, beyond which stimulation effects are considered undesirable.

| Metric | Relevance | <2.5 kHz | 2.5–100 kHz |
| --- | --- | --- | --- |
| E-field brain (peak) | brain stimulation | 16 mA (30 V/m, DBS outside stimulation zone) | 16 mA*f/2.5 kHz (30 V/m*f/2.5 kHz, DBS outside stimulation zone) |
| E-field skin (peak) | skin stimulation | **7 mA (200 V/m, tACS)** | **7 mA\*f/2.5 kHz (200 V/m\*f/2.5kHz, tACS)** |
| total current (peak) | electrode-tissue interface effects | 18 mA (DBS outside sphere) | 18 mA*f/2.5kHz (DBS sphere with frequency scaling) |
| charge/phase (peak) | charge accumulation | 400 mA*f/1 kHz (1.3 mC, tACS) | 400 mA*f/1 kHz (1.3 mC, tACS) |
| brain temperature increase (peak) | brain heating | 14 mA (0.1°C, FDA) | **14 mA (0.1°C, FDA)** |
| skin temperature increase (peak) | skin heating | 100 mA (2°C, FDA) | 100 mA (2°C, FDA) |
| applied voltage (peak-to-peak) | leakage current | 60 V (IEC/ISO 60601-1) | 60 V (IEC/ISO 60601-1) |

We note that these limits do not consider E-field-induced sensations in the skin, since they manifest well before safety becomes relevant. Drawing on available data, TIS currents of at least 7 mA can be applied without exceeding skin field exposures commonplace in tES. Assuming that thresholds scale linearly with frequency above 2.5 kHz, this implies an acceptable current level of >10 mA at 4 kHz and >200 mA at 100 kHz. Rudimentary simulations of 1 μm C-fiber stimulation (relevant for sympathetic neural activation; A- and B-fibers in the skin affect sensation and are not safety relevant) suggest that activation occurs at approximately 1.6 mA, 6 mA, 160 mA, and 1000 mA, for 1 kHz, 2.5 kHz, 4 kHz, and 100 kHz, respectively. Sensations have been reported during stimulation in the low kHz frequency range for currents >2–4 mA^92^. Average current density (ICNIRP 2010 averaging protocol^84^) in the skin beneath the electrode was on the order of 0.1–1.5 mA/cm^2^ per mA of electrode current (electrode size dependent), and 0.008–0.05 mA/cm^2^ per mA of applied current in brain tissue below the electrode. In rodents, unaveraged current densities of up to 32 mA/cm^2^ (500 μA over a 1.6 mm^2^ electrode) at 1 kHz, applied repeatedly in 6 s intervals, have been reportedly used without inducing behavioral AEs^11^.

## Discussion

In light of the sparsity of empirical studies that employ TIS as a neuromodulatory strategy, we endeavored here to synthesize what data does exist regarding the safety of tACS, tDCS, and DBS with respect to thermal effects and unwanted stimulation to identify safe operating conditions for TIS in terms of current intensities, frequencies, electrode placement, and target regions. TIS and tACS are readily compared owing to the close correspondence of the (bio)physical principles at play. In particular, TIS and tACS share a system for transcranial current delivery, permitting careful extrapolation of safety aspects from tACS to TIS with due consideration to differences in operating frequency (within a few tens of Hz for tACS, a few kHz for TIS) and the interferential nature of TIS. On the other hand, DBS, like TIS, aims at selectively activating deep brain structures, providing a qualified reference point for the expected biophysical response to optimally focused TIS. While a complete characterization of TIS safety in humans awaits additional investigation, we can draw upon available research and clinical experience with related technologies to help define the relevant endpoints for monitoring, as regards both safety and effectiveness.

### Physical and Chemical Effects

To date, there have few reports of TIS-associated AEs, all mild to moderate, and with no significant differences between sham and active conditions^11,79,93^. For carrier frequencies <2 kHz and current amplitudes <2 mA, TIS is comparable to tACS in terms of the magnitude and extent of fields and currents, and no physical or chemical damage has been observed for either technique. Furthermore, safety guidelines suggest that safety thresholds for applied current are proportional to frequency above 2.5 kHz. However, empirical data for TIS are lacking, and such extrapolation should be supported by basic safety experiments. Using detailed simulations in a human head model with an input current of 1 mA and an electrode area of 3 cm^2^, we found that skin heating is on the order of a few milliCelsius, while spatial peak current density and field strengths are on the order of 5 A/m^2^ and 30 V/m, respectively. According to the Shannon model for charge-induced tissue damage, this yields comparable exposure values to those of local stimulation methods such as DBS. Furthermore, the electrode materials and conductive gel should be chosen to minimize irritation to the skin (see *<u>tES and AEs</u>*).

### Skin Sensations, Phosphenes, and Ramping

The use of kHz frequencies in TIS considerably increases direct stimulation thresholds in the scalp, where little constructive interference is present, as compared to the lower frequencies characteristic of tACS. Thus, tingling sensations, phosphenes, and unwanted stimulation of facial nerves are improbable for currents ≤2 mA and are unlikely to confound blinding procedures. Existing skin exposure guidelines and the established history of safe use for tES further ensure safety with regards to unwanted facial nerve stimulation, especially when frequency-dependent response attenuation is accounted for. Nevertheless, such sensations must be carefully monitored and human subjects/patients should be questioned regarding their occurrence. It has been observed for TIS in particular that ramping up the current strength rapidly causes transient neural activity^11^, which may cross the threshold for perception. Therefore, care must be taken to increase currents sufficiently slowly; ramping times of a few seconds appear sufficient to avoid such stimulation artifacts (personal correspondence with collaborators). These observations are consistent with the biophysical principles outlined in *Appendix B* and *<u>Computational Characterization and Comparison of Electric Brain Stimulation Modalities</u>*.

### Electrode Shape and Placement

The choice of electrode shape and placement is driven both by the stimulation target and by safety concerns. In general, power deposition and exposure-induced heating are minor, and limited by skin sensations. Nevertheless, practitioners may seek to minimize these quantities by selecting larger electrodes, ensuring high-quality electrical contacts, and by avoiding sharp edges. Employing larger surfaces is practical for remote return electrodes, but may compromise treatment efficacy when applied to electrodes near the target region due to focality degradation (more current overlap in overlying structures), necessitating an analysis of the risk-reward tradeoff when selecting electrode shape and placement. Particular caution is advised when multiple electrodes are placed in close proximity due to the superposition of edge effects. Our analysis of edge and proximity effects also applies in the case of passive electrodes (electrodes that do not provide current), electrodes operated at different frequencies (as required for TIS), and invasively placed electrodes (both sensing and stimulating), which tend to concentrate the local field. Furthermore, the spatial extent of the conductive contact gel must also be considered; seemingly well-separated electrodes may effectively behave as though they were in close proximity if the gel extends sufficiently, while large electrode areas can still result in small contact areas for poor gel coverage. Proper skin and electrode preparation is necessary to mitigate the risk of persistent skin lesions and subjects should be encouraged to report any discomfort to prevent skin irritation^17^. Finally, in cases where a single electrode is used as the common return electrode for multiple TIS current channels, it is important to consider that the total power deposition – and similarly the total temperature increase – is equal to the summed combination from all channels (incoherent field superposition).

### Short- and Long-Term Neuromodulatory Effects

At a standard tES current amplitude of approximately 1 mA per channel, direct suprathreshold TIS is not expected from either the high-frequency carrier or the low-frequency modulation. Instead, neuromodulation occurs in the subthreshold regime via membrane polarization and synchronization of network activity. Conservatively, AEs associated with tACS (see *<u>tES and AEs</u>*) should be monitored for when applying TIS. However, as kHz frequencies are unlikely to evoke stimulation, and since the low frequency modulation must first undergo a rectification and/or mixing process, the actual stimulation efficiency of TIS is likely lower than that of tACS for the same exposure magnitude. Furthermore, as TIS aims to produce localized stimulation in deep brain structures, DBS-like AEs are also possible (see Table A.4, *Appendix A*). Yet, because local DBS brain exposure strengths are generally two orders of magnitude higher than those of TIS, and since (unlike TIS) DBS is applied chronically for months or years, DBS-associated AEs are presumably less likely for TIS.

The analogy between TIS and tACS/DBS has several limitations. In particular, direct comparisons between the E-field magnitude of low-frequency stimulation and the TIS modulation magnitude are precluded by the unknown efficacy of signal rectification and/or nonlinear frequency mixing by neurons. Comparing high-frequency field exposure strengths in TIS to the field strengths in tACS/DBS is also not appropriate given the discrepancy in frequencies and attendant differences in the neural response.

Little is known about the long-term effects of TIS. Particularly for modulation frequencies in the alpha to theta band, attention must be paid to the possibility of alterations in various cognitive functions. Following the logic of Klink et al.^4^, if cognitive enhancements are possible with tACS, then AEs in the domain of cognition should also be possible. Furthermore, as long term TIS applied in regular sessions over a period of weeks or months may lead to plasticity-dependent changes, treatment cessation could result in withdrawal symptoms, the occurrence of which should be monitored.

### Planning, Monitoring, and Reporting

In TIS, the added complexity of selectively targeting deep structures using multiple, interacting currents/fields requires proper treatment planning and optimization, which is generally undertaken using computational modeling. Simulations are strongly recommended, both as a means of improving focality, and as a source of exposure information to guide safety assessments. It remains to be seen whether and how individualization of treatment parameters to accommodate inter-subject anatomical variations affect treatment outcomes. Should such differences prove significant, subject-specific treatment planning will be required for the design of optimal stimulation configurations. Due to the novelty of TIS and the importance of adhering to safety protocols, the planning and delivery of TIS should be executed by trained personnel. We refer the reader to a recent review paper by the International Federation of Clinical Neurophysiology (IFCN) summarizing the requirements for training^94^.

### Risk-Benefit Analysis

Similar to other brain stimulation approaches, a thorough risk-benefit analysis must undergird both clinical and research applications. For example, DBS is usually applied in refractory diseases where the considerable risks of surgery are worth the anticipated benefits. In the case of TIS, the risk-benefit balance of TIS is likely to be favorable in most circumstances, since the potential for therapeutic benefits are appreciable, with minimal risk of AEs. Particular caution must be exercised when targeting deep brain structures in view of their central role in regulating core physiological functions (e.g., heartbeat, breathing, thermoregulation), and the potential to trigger anxiety or other severe responses. In particular, structures such as the amygdala (emotion/fear processing) or regions in the brain stem should be avoided. Indications regarding critical brain structures might be derived from observed AEs related to DBS (Table A.4, *<u>Appendix A</u>*). As with tES, care is required in identifying patient populations with heightened susceptibility to certain AEs (e.g., predisposition for manic episodes, prior history/family history of neurologic/psychiatric disorders or seizures). Despite one report of seizure, many thousands of sessions of tES have been performed without incident. Thus, accurate patient history and seizure propensity should be taken into account before tES treatments, but do not necessarily preclude subjects from tES treatment. Particular caution must also be exercised when exposing populations such as infants, children and/or adolescents, whose brains are still developing.

### Mechanistic Pathway and TI Planning Optimization Metric

The current lack of a clear mechanism(s) of action for TIS complicates safety assessments and treatment optimization, in part by frustrating efforts to establish a suitable exposure metric that relates to stimulation efficacy and selectivity. For example, the observed shift of the TIS focus towards the weaker current source^11^ agrees with predictions based on the modulation envelope magnitude distribution. In contrast, a stimulation mechanism based on 2^nd^ order frequency mixing would predict an effect size that scales with the product of the local exposure magnitudes of the two sources and therefore, despite support from electrophysiological data, cannot explain steering. Furthermore, in highly structured tissues (e.g., layer 5 cortical pyramidal cells, hippocampal complexes, collinear axonal projections), the preferred field orientation of a neural population likely requires the consideration of projected field quantities, rather than the full field magnitude. Activating function (AF) concepts, as suggested by Mirzakhalili et al., also imply that field heterogeneity, rather than field magnitude alone, must be considered^95^. Other important factors include the influence of ongoing neural activity, and differences between brain regions in terms of susceptibility to stimulation. Therefore, dedicated experiments designed to explore the mechanisms of action of TIS are needed.

As a corollary to the above discussion, the modeling work presented here is limited by the assumption of the homogeneity of brain tissue biophysical properties. Thus, cell type-specific properties including membrane polarizability and threshold potential are not considered in the predicted response to applied E-fields or TI amplitude modulation. These difficulties are further compounded in the absence of knowledge concerning TIS mechanisms, which may exert differential effects depending on ongoing neural dynamics or subpopulation characteristics. The availability of such information is limited, though attempts have been made to aggregate knowledge of the neural properties of certain populations (e.g., in the hippocampus)^96^.

### Physical Limits on Targeting

An intriguing benefit of TIS is the potential for non-invasive steering of the focus^11^. However, the physics of EM fields, in particular the impedances of different pathways to current flow, introduces complexities that may either inhibit or facilitate targeting. For example, longer paths along highly conductive media (e.g., CSF) may have less overall impedance than shorter paths through highly resistive media (e.g., cortical bone). Thus, geometric length is not identical to effective electrical length. Current may flow through the scalp to locations where the skull offers less resistance or may preferentially follow paths that maximize flow through highly conductive CSF, while reducing the length of passage through brain tissue (“current shunting”). Similarly, brain tissue, especially white matter, is anisotropic, and conductivity along fiber orientations can be up to an order of magnitude higher than in the transverse direction. Computational modeling allows for the consideration of all of these factors during optimization and treatment planning.

### Implants

Conductive implants (e.g., invasive stimulation and/or recording electrodes) pose a safety concern, as they can lead to significant local field increases, affecting both the exposure magnitudes and precision of targeting.

### Training

Specialized training for personnel who administer TIS is recommended (see for example Fried et al.)^94^.

### Limitations

In general, it should be noted that we cannot rule out bias in the selection of the tACS, tDCS, and DBS literature considered here, as our search focused on recent reviews. In addition, literature reporting negative results was generally not included (and is less likely to have been published). We also note the following limitations affecting our analysis of the simulation data reported in the preceding sections:

- Since the exact mechanisms of TI exposure-related neuromodulation have not been clarified, there remains a possibility that exposure metrics other than those described here are relevant to safety.
- Additional simulations exhausting the plausible stimulation configurations used in tACS, tDCS, TIS and DBS are required. Moreover, an analysis of uncertainty is needed to ascertain the confidence intervals associated with our model-based predictions.
- Little experimental data exists for brain stimulation at high frequency, necessitating physics-based extrapolations from stimulation at other frequencies.
- Proposed limits for exposure-related quantities apply to adult heads, and should not be used in investigations that include pediatric subjects, or subjects with anatomical anomalies (e.g., skull fractures, brain lesions, implanted electrodes).

## Conclusion and Outlook

TIS promises to unlock new opportunities for non-invasive, targeted, deep brain stimulation. However, as an emerging technique, considerable effort is still needed to establish a foundation of knowledge to support future developments. In particular, guidance regarding exposure limits and other safety concerns, necessary for obtaining regulatory approvals and conducting research, is lacking. To this end, we drew upon the already available data and expertise in electrical neuromodulation to inform a preliminary analysis of TIS exposure safety. To compensate for gaps in the TIS literature, biophysical arguments were used to translate knowledge from common tES and DBS modalities to TIS, considering a wide range of safety-relevant interactions. First, we reviewed and summarized the biophysical mechanisms, potential safety concerns, and reported AEs of tES and DBS. Next, we employed dosimetric and biophysical modeling to support comparisons across different electrical neurostimulation modalities. TIS opens the door to new applications involving the selective, non-invasive stimulation of deep brain regions. However, such regions play important roles in regulating essential physiological and emotional processes, warranting special caution in the application of TIS.

## Disclosures

Esra Neufeld is a minority shareholder and a board member of TI Solutions AG.

Niels Kuster is a shareholder of NF Technology Holdings AG, which is a minority shareholder of TI Solutions AG. He is also a board member of TI Solutions AG.

Sabine Regel is the CEO of TI Solutions AG.

## Acknowledgements

The authors thank Profs. Ed Boyden, Nir Grossman and Alvaro Pascual-Leone for their valuable input.

## Acronyms

AC: Alternating current
A-CalCAP: California computerized assessment package
ACC: Anterior cingulate cortex
AD: Alzheimer’s disease
AE: Adverse event
AF: Activating function
ALIC: Anterior limb of the internal capsule
ANT: Anterior nucleus of the thalamus
AP: Action potential
AUC: Area under the curve
BNST: Bed nucleus of the stria terminalis
CB: Conduction blocking
CMN: Centromedian nucleus of the thalamus
CM-PFc: Centromedian-parafascicular thalamic complex
CN: Caudate nucleus
CNS: Central nervous system
CSF: Cerebral spinal fluid
cZI: Caudal zona incerta
DBS: Deep brain stimulation
DC: Direct current
DTI: Diffusion tensor imaging
ECT: Electroconvulsive therapy
EM: Electromagnetic
ET: Essential tremor
FDA: U.S. food and drug administration
GABAB: Gamma-aminobutyric acid
GP: Globus pallidus
GPe: External globus pallidus
GPi: Internal globus pallidus
HD-tES: High-density tES
Hipp: Hippocampus
IPG: Implanted pulse generator
ISI: Interspike interval
ITP: Inferior thalamic peduncle
LTD: Long-term depression
LTP: Long-term potentiation
MAUDE: Manufacturer and user facility device experience
MD: Major depression
MFB: Medial forebrain bundle
mNSS: Modified neurological severity score
MoCA: Montreal cognitive assessment
MRI: Magnetic resonance imaging
NAcc: Nucleus accumbens
NBM: Nucleus basalis of Meynert
NIBS: Non-invasive brain stimulation
NMDA: *N*-methyl-D-aspartate
NSE: Neuron-specific enolase
OCD: Obsessive-compulsive disorder
PAG: Periaqueductal grey matter
PBE: Pennes Bioheat Equation
PD: Parkinson’s disease
pHr: Posterior hypothalamic region
PMA: Premarket approval
PMS: Postmarket surveillance
PNS: Peripheral nervous system
PPN: Pedunculopontine nucleus
PPT: Purdue pegboard test
PSA: Posterior subthalamic area
PSC: Postsynaptic current
PVG: Periventricular grey matter
RF: Radiofrequency
RMS: Root mean square
SAR: Specific absorption rate
SAS: Self-assessment scale
SAT: Subcutaneous fat
SD: Standard deviation
SCC: Subcallosal cingulate cortex
SCG: Subgenual cingulate gyrus (=BA25=Cg25)
SNpr: Substantia nigra pars reticulata
SNR: Signal-to-noise ratio
STN: Subthalamic nucleus
tACS: Transcranial alternating current stimulation
tDCS: Transcranial direct current stimulation
tES: Transcranial electrical stimulation
TI: Temporal interference
TIS: Temporal interference stimulation
TMS: Transcranial magnetic stimulation
TPLC: Total product life cycle
VAMS-R: Visual analog mood scale
VC: Ventral capsule
VIM: Ventral intermedius nucleus of the thalamus
VO: Ventral oralis nucleus of the thalamus
VPL: Ventral posterior lateral nucleus
VPM: Ventral posterior medial nucleus
VS: Ventral striatum
VTA: Ventral tegmental area
ZI: Zona incerta

## Appendix A

**Table A.1:**
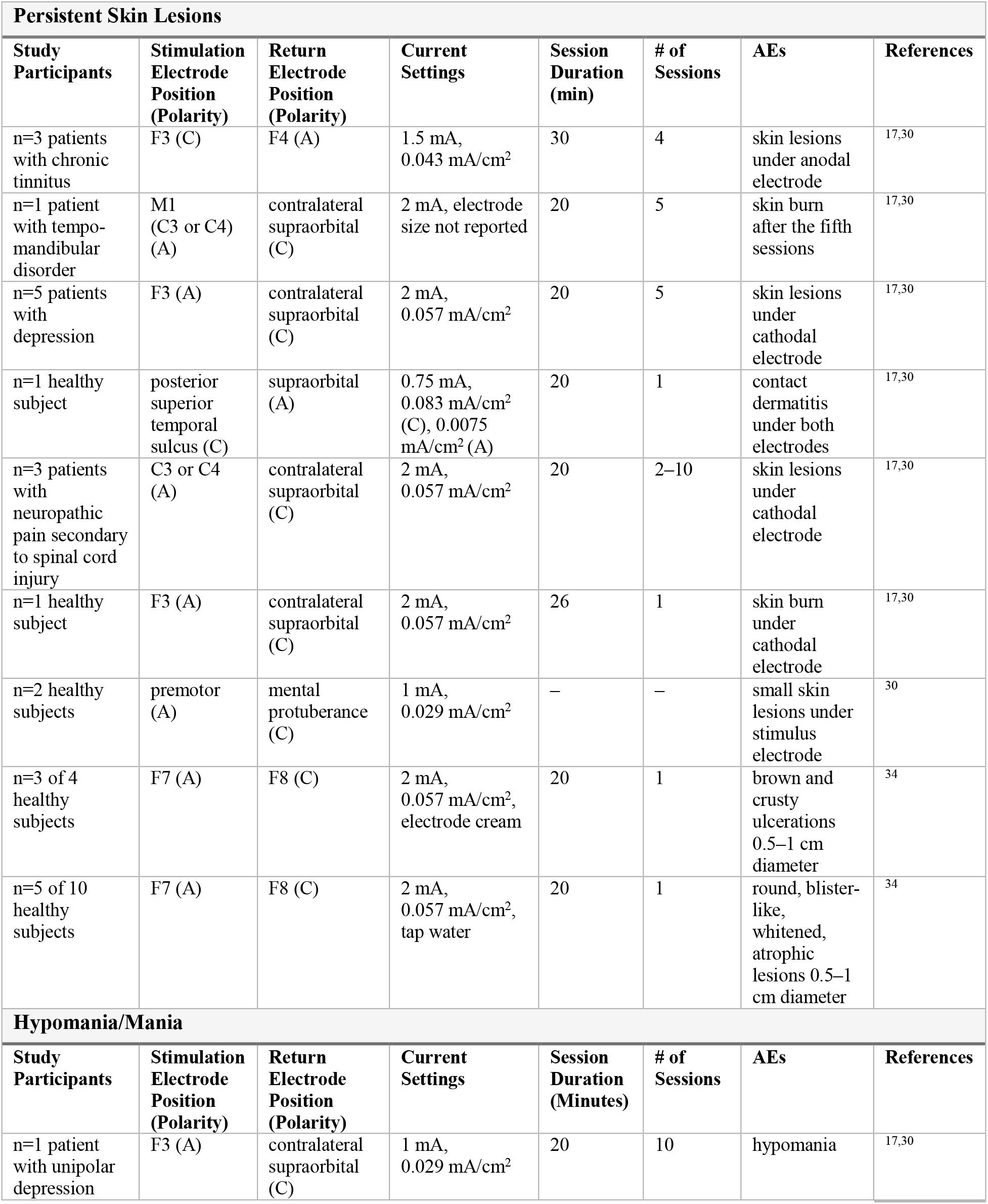
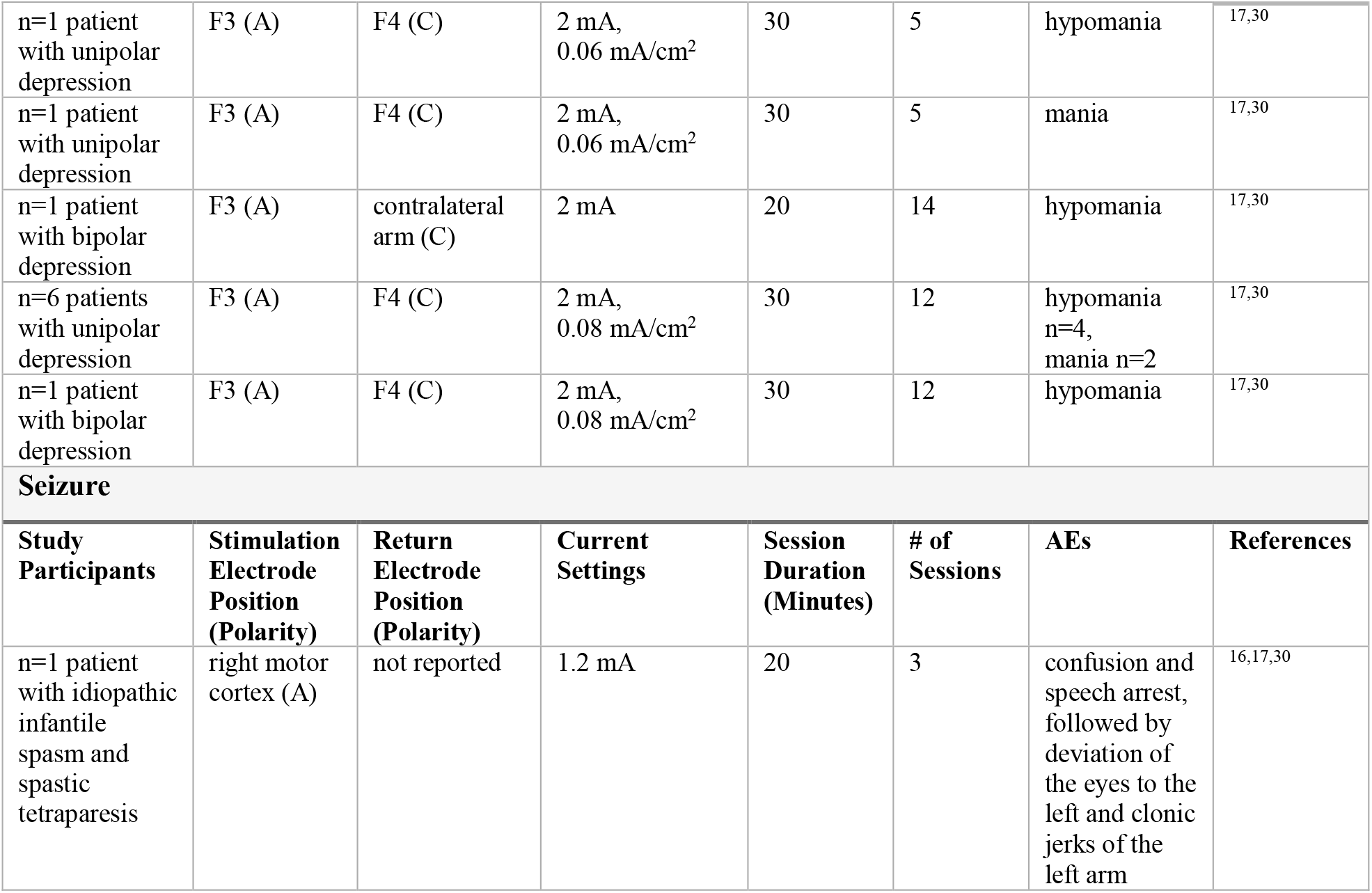
tES and AEs. Summary of moderate to severe AEs reported for tDCS with associated stimulation parameters. A: anode, C: cathode.

| Persistent Skin Lesions |  |  |  |  |  |  |  |
| --- | --- | --- | --- | --- | --- | --- | --- |
| Study Participants | Stimulation Electrode Position (Polarity) | Return Electrode Position (Polarity) | Current Settings | Session Duration (min) | # of Sessions | AEs | References |
| n=3 patients with chronic tinnitus | F3 (C) | F4 (A) | 1.5 mA, 0.043 mA/cm<sup>2</sup> | 30 | 4 | skin lesions under anodal electrode | 17,30 |
| n=1 patient with tempo-mandibular disorder | M1 (C3 or C4) (A) | contralateral supraorbital (C) | 2 mA, electrode size not reported | 20 | 5 | skin burn after the fifth sessions | 17,30 |
| n=5 patients with depression | F3 (A) | contralateral supraorbital (C) | 2 mA, 0.057 mA/cm<sup>2</sup> | 20 | 5 | skin lesions under cathodal electrode | 17,30 |
| n=1 healthy subject | posterior superior temporal sulcus (C) | supraorbital (A) | 0.75 mA, 0.083 mA/cm<sup>2</sup> (C), 0.0075 mA/cm<sup>2</sup> (A) | 20 | 1 | contact dermatitis under both electrodes | 17,30 |
| n=3 patients with neuropathic pain secondary to spinal cord injury | C3 or C4 (A) | contralateral supraorbital (C) | 2 mA, 0.057 mA/cm<sup>2</sup> | 20 | 2–10 | skin lesions under cathodal electrode | 17,30 |
| n=1 healthy subject | F3 (A) | contralateral supraorbital (C) | 2 mA, 0.057 mA/cm<sup>2</sup> | 26 | 1 | skin burn under cathodal electrode | 17,30 |
| n=2 healthy subjects | premotor (A) | mental protuberance (C) | 1 mA, 0.029 mA/cm<sup>2</sup> | — | — | small skin lesions under stimulus electrode | 30 |
| n=3 of 4 healthy subjects | F7 (A) | F8 (C) | 2 mA, 0.057 mA/cm<sup>2</sup>, electrode cream | 20 | 1 | brown and crusty ulcerations 0.5–1 cm diameter | 34 |
| n=5 of 10 healthy subjects | F7 (A) | F8 (C) | 2 mA, 0.057 mA/cm<sup>2</sup>, tap water | 20 | 1 | round, blister-like, whitened, atrophic lesions 0.5–1 cm diameter | 34 |
| Hypomania/Mania |  |  |  |  |  |  |  |
| Study Participants | Stimulation Electrode Position (Polarity) | Return Electrode Position (Polarity) | Current Settings | Session Duration (Minutes) | # of Sessions | AEs | References |
| n=1 patient with unipolar depression | F3 (A) | contralateral supraorbital (C) | 1 mA, 0.029 mA/cm<sup>2</sup> | 20 | 10 | hypomania | 17,30 |
| n=1 patient with unipolar depression | F3 (A) | F4 (C) | 2 mA, 0.06 mA/cm<sup>2</sup> | 30 | 5 | hypomania | 17,30 |
| n=1 patient with unipolar depression | F3 (A) | F4 (C) | 2 mA, 0.06 mA/cm<sup>2</sup> | 30 | 5 | mania | 17,30 |
| n=1 patient with bipolar depression | F3 (A) | contralateral arm (C) | 2 mA | 20 | 14 | hypomania | 17,30 |
| n=6 patients with unipolar depression | F3 (A) | F4 (C) | 2 mA, 0.08 mA/cm<sup>2</sup> | 30 | 12 | hypomania n=4, mania n=2 | 17,30 |
| n=1 patient with bipolar depression | F3 (A) | F4 (C) | 2 mA, 0.08 mA/cm<sup>2</sup> | 30 | 12 | hypomania | 17,30 |
| **Seizure** |  |  |  |  |  |  |  |
| **Study Participants** | **Stimulation Electrode Position (Polarity)** | **Return Electrode Position (Polarity)** | **Current Settings** | **Session Duration (Minutes)** | **# of Sessions** | **AEs** | **References** |
| n=1 patient with idiopathic infantile spasm and spastic tetraparesis | right motor cortex (A) | not reported | 1.2 mA | 20 | 3 | confusion and speech arrest, followed by deviation of the eyes to the left and clonic jerks of the left arm | 16,17,30 |

**Table A.2:** DBS Targets. List of DBS targets by clinical indication in alphabetical order. *well established, **not widely implemented, still under investigation, ***very limited investigation/largely hypothetical^47^. Note that for drug-resistant epilepsy, the number of brain targets may be considerably larger as stimulation is also applied to the ictal onset zone, i.e., the location causing the seizures, which varies widely across patients. To date, only DBS targeting the ANT has been recognized for epilepsy by the FDA^64^. See the list of *<u>Acronyms</u>* for DBS target names.

| Disease | Circuit<sup>6</sup> | DBS Target | References |
| --- | --- | --- | --- |
| addiction/<br>substance abuse | reward | ALIC<br>NAcc***<br>STN | 3,57,97,98 |
| aggressive<br>behavior/self-harm | n/a | amyg<br>GPi<br>pHr | 3,99 |
| Alzheimer's disease | cognitive and<br>memory circuits | fornix***<br>NBM<br>VC/VS<br>ventromed. prefrontal<br>cortex*** | 5,47,100–102 |
| anorexia nervosa | reward and mood | BNST<br>MFB<br>NAcc***<br>SCC<br>SCG<br>VC/VS | 3,99,103–105 |
| chronic pain | sensory systems and<br>interoceptive<br>awareness | ACC**<br>ALIC<br>CM-PFc<br>PAG/PVG**<br>VPL/VPM**<br>VS | 57 |
| cluster headache | n/a | post. hypothalamus**<br>floor of 3rd ventricle<br>VTA | 47,106 |
| dystonia | motor | GPi*<br>thalamus<br>STN<br>subthalamic area | 107 |
| epilepsy | various | amyg<br>ANT<br>cerebellum<br>CMN<br>CM-PFc<br>Hipp**<br>NAcc<br>SNpr<br>STN<br>cZI | 5,57,103,108,109 |
| essential tremor | motor | VIM*<br>PSA/cZI<br>ZI** | 47,57,110 |
| Gilles de la Tourette syndrome | motor and limbic | ALIC<br>CMN<br>CM-PFc<br>CN<br>GPe<br>GPi**<br>NAcc<br>STN<br>thalamus<br>VC/VS<br>VO | 3,57,99,111 |
| Huntington's disease | n/a | GPe<br>GPi***<br>STN | 47,112 |
| major depression | limbic | ALIC<br>BNST<br>ITP<br>LHb<br>MFB<br>NAcc**<br>SCC<br>SCG**<br>VC/VS | 3,99 |
| multiple sclerosis<br>Tremor | n/a | STN<br>thalamus/ventrolat. thalamus<br>VC<br>VIM<br>VIM/VO<br>VO/ZI<br>ZI | 113–115 |
| obesity | n/a | hypothalamus (lateral,<br>ventromedial)<br>NAcc*** | 57,99,105,116 |
| obsessive compulsive disorder | motor and limbic | ALIC**<br>BNST<br>CN<br>GPi<br>ITP<br>NAcc<br>Thalamus (medial)**<br>STN<br>VC/VS | 3,57,73,99,117–120 |
| Parkinson's disease | motor | GPi*<br>NBM<br>PPN**<br>STN*<br>VIM<br>ZI** | 47,57,101,102,121,122 |
| tinnitus | auditory | ACC<br>CN<br>GPi<br>STN<br>VIM | 123 |

**Table A.3:** DBS Stimulation Parameters. List of DBS stimulation parameters by clinical indication in alphabetical order. Note that data regarding stimulation parameters was not available for all indications. In cases where mean values and standard deviations (SD) were provided, the range was estimated as +/- 2 SD from the mean.

| Disease | Circuit<sup>6</sup> | Frequency | Voltage or Current Amplitude | Pulse Width | References |
| --- | --- | --- | --- | --- | --- |
| addiction/substance abuse | reward | 130.0–185.0 Hz | 1.5–7.0 V | 90.0–240.0 μs | 97,98 |
| aggressive behavior/self-harm | n/a | – | – | – | – |
| Alzheimer's disease | cognitive and memory circuits | 5.0–130.0 Hz | 1.0–5.0 V | 60.0–210.0 μs | 5,100,102 |
| anorexia nervosa | reward and mood | 130.0–185.0 Hz | 2.5–7.0 V<br>5 μA | 90.0–210.0 μs | 105 |
| chronic pain | sensory systems and interoceptive awareness | – | – | – | – |
| cluster headache | n/a | 130.0–185.0 Hz | 0.7–3.7 V | 60.0–120.0 μs | 106 |
| dystonia | motor | GPi:<br>95.0–190.0 Hz<br>thalamus:<br>120.0–180.0 Hz<br>STN:<br>65.0–220.0 Hz<br>subthalamic area:<br>100.0–180.0 Hz | GPi:<br>1.0–5.0 V<br>thalamus:<br>1.7–3.3 V<br>STN:<br>1.25–4.0 V<br>subthalamic area:<br>1.8–2.6 V | GPi:<br>30.0–220.0 μs<br>thalamus:<br>45.0–160.0 μs<br>STN:<br>15.0–155.0 μs<br>subthalamic area:<br>55.0–110.0 μs | 107,124 |
| | | across targets:<br>65.0–220.0 Hz | across targets:<br>1.0–5.0 V | across targets:<br>15.0–220.0 μs | |
| epilepsy | various | 2.0–333.0 Hz | 0.5–14.0 V<br>0.2–12.0 mA | 40.0–520.0 μs | 108,109 |
| essential tremor | motor | – | 0.4–6.2 V | 60.0–140.0 μs | 110 |
| Gilles de la<br>Tourette<br>syndrome | motor and<br>limbic | 102.0–170.0 Hz | 3.1–6.4 V | 108.0–210.0 μs | 125 |
| Huntington's<br>disease | n/a | across targets:<br>40.0–180.0 Hz | GPI:<br>0.5–3.0 V<br>STN:<br>2.0–5.0 V<br>across targets:<br>0.5–5.0 V | GPI:<br>90.0 μs<br>STN:<br>60.0 μs<br>across targets:<br>60.0 μs and 90.0<br>μs | 112 |
| major depression | limbic | 5.0–185.0 Hz | 1.5–10.5 V<br>2.0–8.0 mA | 30.0–450.0 μs | 126,127 |
| Multiple Sclerosis<br>tremor | n/a | 90.0–160.0 Hz | 1.5 V–4.8 V | 60.0–120.0 μs | 114 |
| obesity | n/a | 8.0–185.0 Hz | 1.0–7.0 V<br>2.0–3.75 mA | 60.0–210.0 μs | 105,116 |
| obsessive<br>compulsive<br>disorder | motor and<br>limbic | 85.0–150.0 Hz | 2.5–10.5 V | 60.0–450.0 μs | 117,120 |
| Parkinson's<br>disease | motor | 20.0–185.0 Hz<br>range causing<br>AEs:<br>50.0–185.0 Hz | 1.0–3.85 V<br>range causing<br>AEs:<br>>2.04 V<br>1.3–3.4 mA | 60.0–120.0 μs<br>– | 70,72,102,122,125 |
| tinnitus | auditory | 150.0–185.0 Hz<br>and<br>6 Hz bursts (each<br>burst = 5 spikes<br>at 500 Hz, 1 ms<br>pulse width) | ≤10.0 V<br>1.4 mA | 60.0–180.0 μs | 123 |

**Table A.4:**
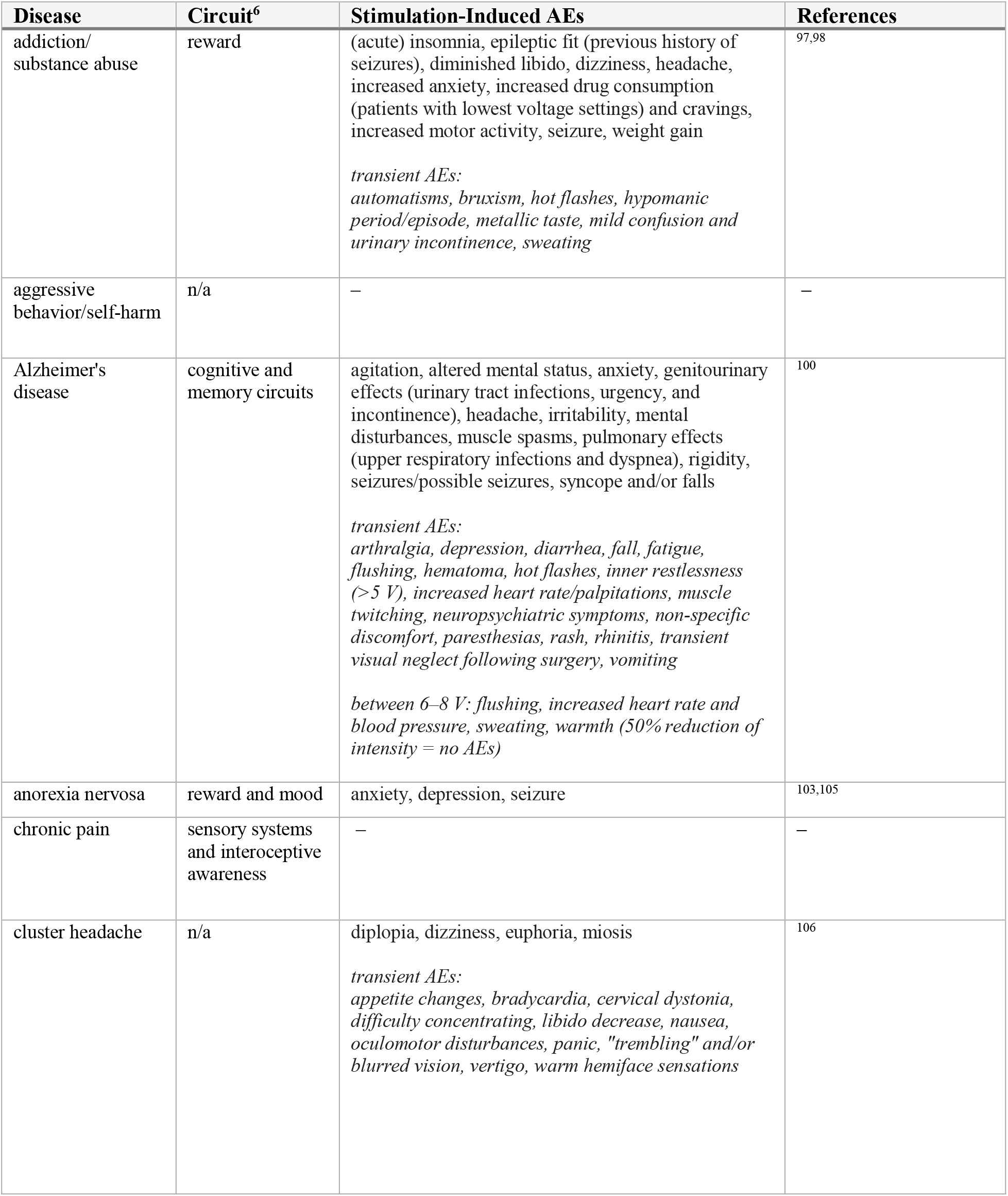
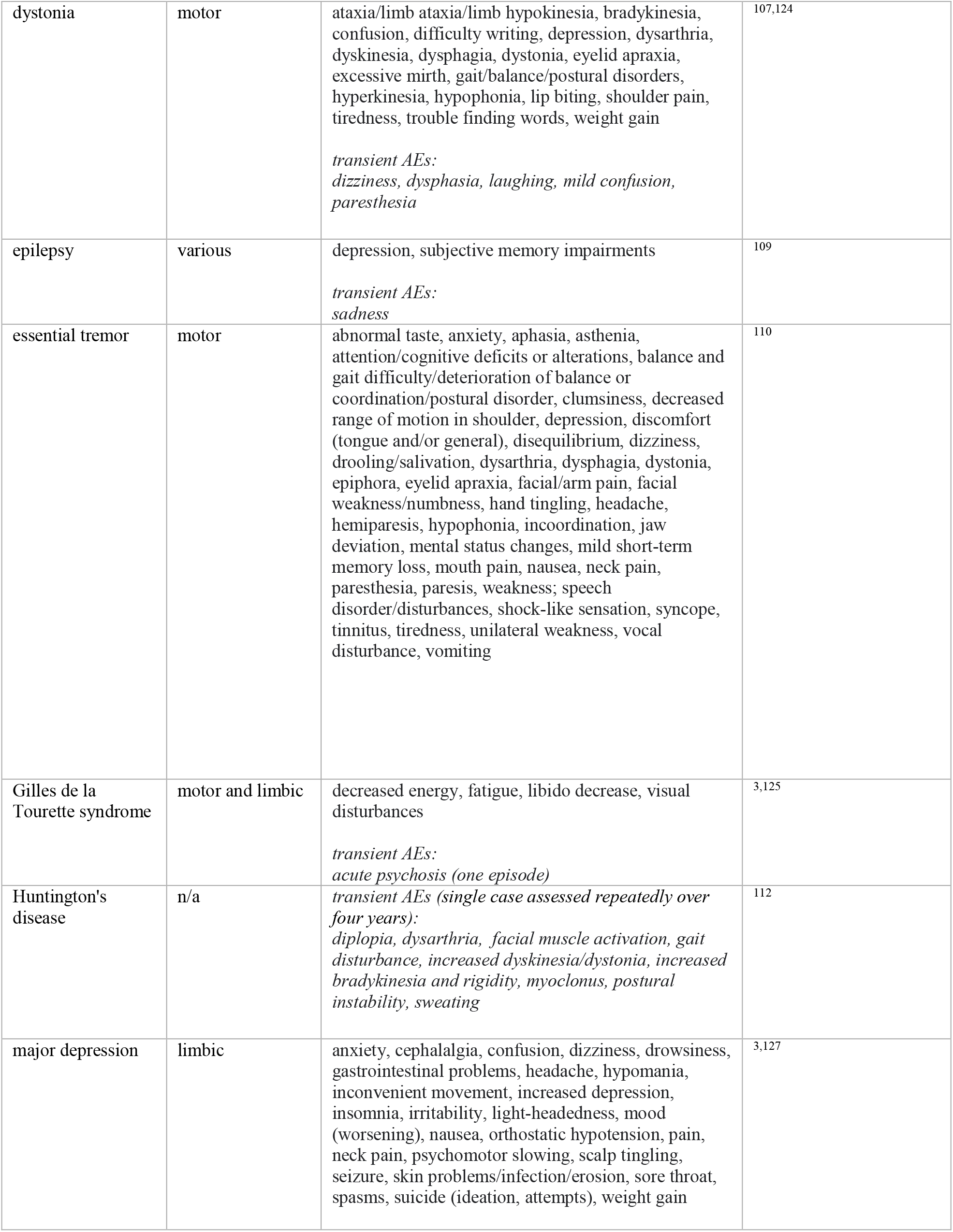
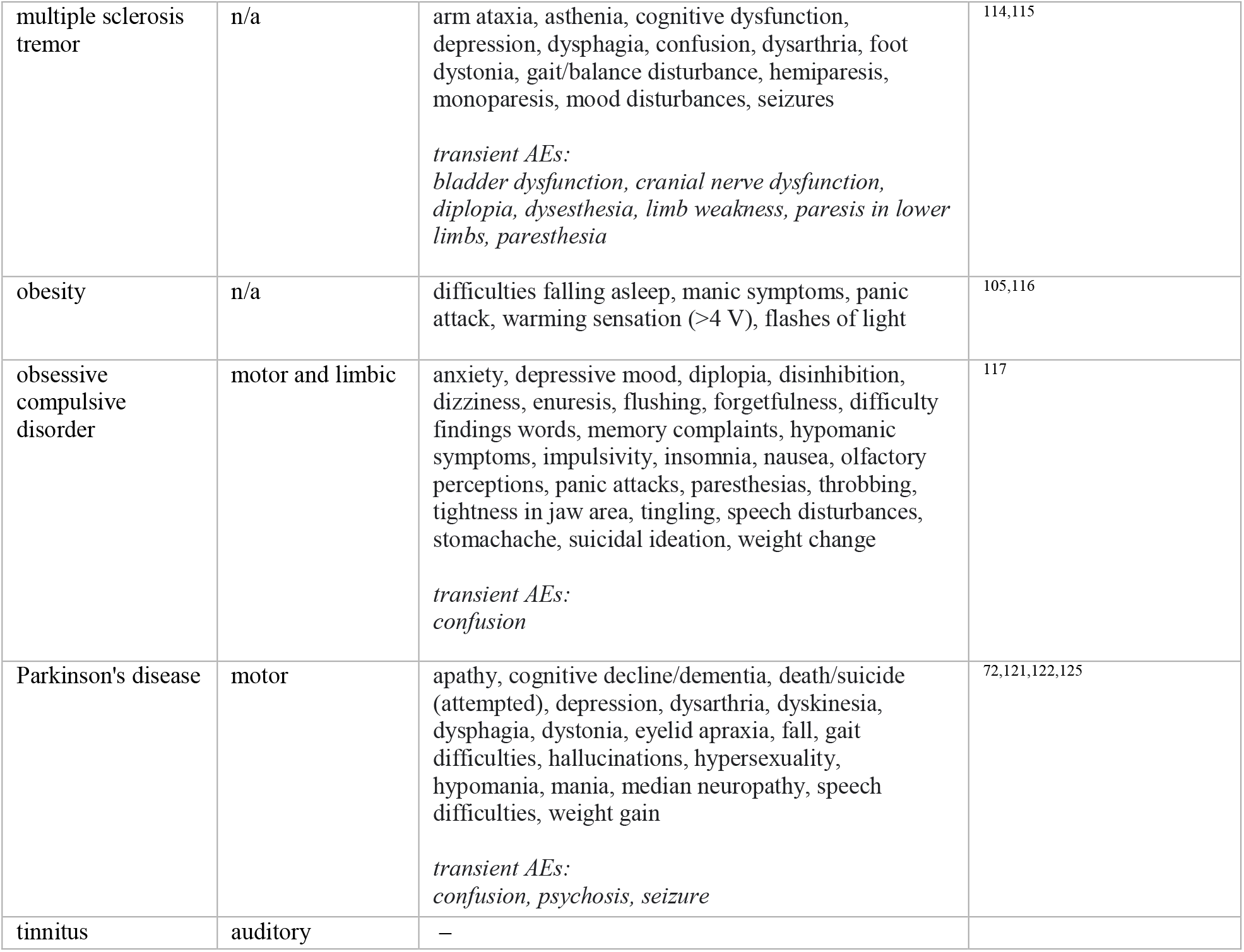
DBS and AEs. List of DBS-induced AEs by clinical indication in alphabetical order. AEs explicitly related to surgery or device failure were not considered. AEs were not distinguished by severity.

## Appendix B

### (Bio)physics of Electrical Stimulation

In this appendix, we begin by reviewing how E-fields interact physically with neurons and highlight relevant exposure metrics to enable comparisons across electrical stimulation regimes. Subsequently, the resulting impacts on neural activity in the sub- and suprathreshold regimes are discussed, distinguishing short- and long-term effects, before reviewing the basic principles of specific therapeutic neurostimulation approaches. While our primary focus is the electrophysiological impact of electrical neurostimulation, other aspects are also discussed in view of potential AEs. We present this information here to build context for comparisons between TIS and tES/DBS, since relating these neuromodulation modalities necessitates an understanding of their mechanisms of action in the brain and associated safety concerns.

#### Biophysical Fundamentals of EM-Neuron Interactions

Low frequency E-fields, typical of neurostimulation applications, interact with neurons via several distinct mechanisms^128^. In general, these mechanisms hinge on the de/hyperpolarization of neural membranes in the presence of EM fields (see Figure B.1). At dendritic and axonal terminals, membrane polarization can be initiated by the tangential E-field component, *E*_*tan*_, which drives axial currents, resulting in a build-up of charge. Alternatively, in non-terminal compartments, charge may accumulate as a consequence of heterogeneous *E*_*tan*_, causing an imbalance between incoming and outgoing axial currents. For long membrane tracts in the resting state, the rate of change of the membrane potential in response to an applied field is given by −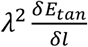 where *λ* is the membrane space constant. This term is referred to as the activating function (AF) and is used to predict the location(s) at which stimulation is likely to occur, as well as the stimulation threshold. A large AF can result from a spatially heterogeneous field or from a rapid change in fiber orientation. A special case of field heterogeneity occurs when fibers cross an interface with dielectric contrast such as at the boundary between tissues with differing electrical conductivities. As shown by Miranda et al., the amplitude of these local E-field discontinuities across the interface is given by 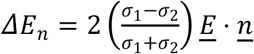, where *<u>E</u>* is the applied field, *σ*_1_ and *σ*_2_ are the conductivities of the two tissues, and *<u>n</u>* is the normal vector at the interface^128^. Tissue heterogeneity-induced gradients most strongly affect axons perpendicular to the interface, inducing a membrane depolarization of *λΔE*_*n*_/2. When the depolarization at neurite terminals is sufficiently large to initiate an AP, this is referred to as end-node stimulation. In contrast, stimulation induced along a fiber tract by any means (tissue heterogeneity or large E-field gradients) is referred to as central-node stimulation.

**Figure B.1.**
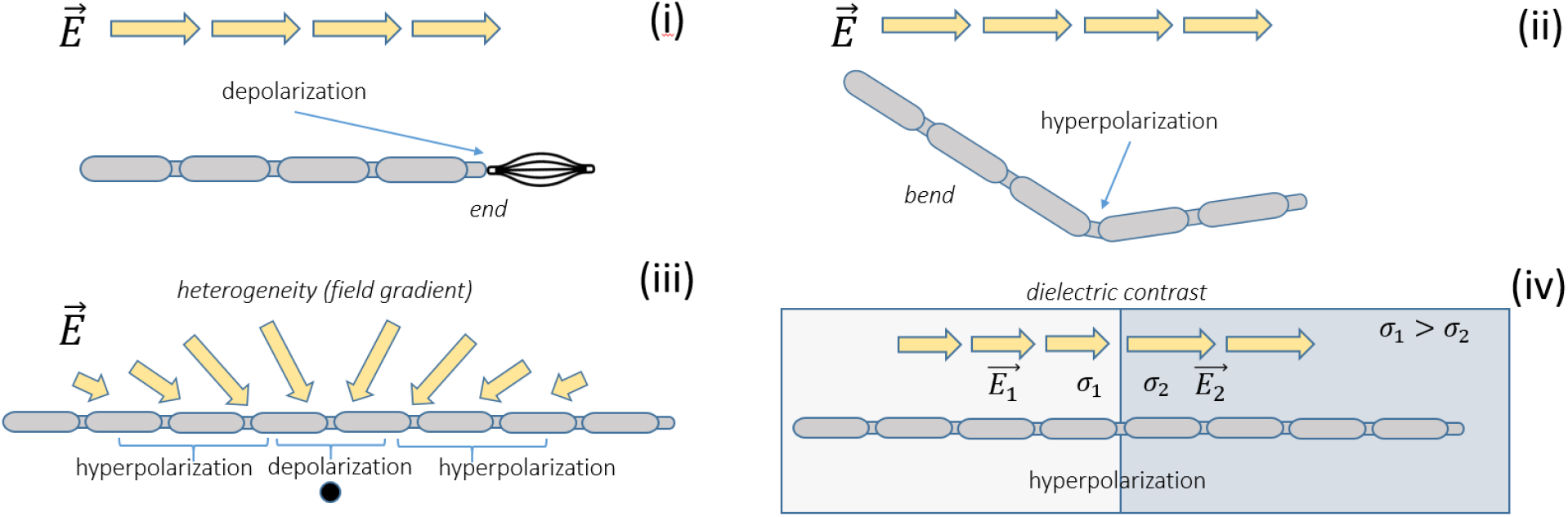
Illustration of different conditions for electric stimulation by: (i) large E-field intensities along the fiber and in proximity to axon terminals (end-node stimulation), (ii) axonal curvature within a homogeneous field, inducing a large E-field gradient along the trajectory, (iii) large E-field gradients along the axon (central-node stimulation), and (iv) local E-field gradients induced at interfaces between tissues with different electrical conductivities.

The relative contributions of these two membrane polarization mechanisms (tangential fields at endings and spatially heterogeneous tangential fields along fibers) are dependent on the modality of brain stimulation. In typical tES setups, where large electrodes (≥ 5 cm^2^) are used, the E-field directly below the electrode is relatively homogeneous, and any field heterogeneity is primarily a consequence of cortical gyrification. Therefore, while sub-millimeter neural somas are exposed to relatively small E-field gradients, neurite termination may cause significant membrane depolarization in dendrites (and through the dendritic arbor, soma polarization). Tissue heterogeneity-related contributions may be relevant for axons with high curvature, for those passing near to CSF, and for those crossing between brain regions with dielectric contrast (e.g., grey to white matter). The situation is reversed when considering DBS stimulation, where the stimulation is applied by small, millimeter-sized electrodes in direct contact with tissue. In this case, axons and somata are exposed to large local E-field gradients within relatively homogeneous regions of tissues (see also *<u>Computational Characterization and Comparison of Electric Brain Stimulation Modalities</u>*).

It is important to note that the degree of exposure-induced membrane depolarization and associated stimulation thresholds are frequency dependent. Thus, low frequency tACS and high frequency TIS carrier signals have different stimulation thresholds, even when exposure strength is matched. For alternating current (AC) exposure at frequencies with periods that are short or comparable with respect to membrane time constants (on the order of 3 ms for unmyelinated C-fibers and 0.3 ms for myelinated fibers), the membrane capacitance results in low-pass filtering, such that polarizability is inversely proportional to frequency. This is reflected in the linearly frequency-dependent exposure thresholds in exposure safety guidelines, such as ^129^. At sufficiently high frequencies, the frequency-dependence of dielectric properties must also be considered when determining *in vivo* exposure conditions.

#### Subthreshold Electrophysiological Impact

##### Immediate Effects

Applied subthreshold E-fields can de/hyperpolarize neurons in direct proportion to the applied field^130^, biasing the timing or probability of spike firing, as smaller/larger membrane potential fluctuations are required to reach the AP threshold. Exogenous E-fields are theorized to modulate neural activity differently depending on the strength of the applied field and the intended outcome. In particular, direct current (DC) anodal stimulation tends to increase neural spiking, while cathodal stimulation tends to suppress it^131,132^, though the magnitude of the effect depends on the neural morphology and its relative orientation to the E-field^130^. Furthermore, second order effects may also play a role, namely membrane polarization of opposite sign in stimulation-adjacent regions as a consequence of local conservation of charge^133^.

For weak applied fields (<1 V/m), exogenous polarization interacts with ongoing dynamics to bias spike times, and theoretically enhance the signal-to-noise ratio (SNR) of the neural code^134^. This effect is known as “stochastic resonance”, though ongoing network activity makes quantification challenging given available techniques^2^. At higher field strengths, network entrainment, wherein applied fields enhance ongoing oscillations (or overcome native brain rhythms to induce new oscillations), is possible^135–137^, though at the expense of increased current requirements^138^. Uniform AC fields have been shown to increase firing time coherence with the applied field oscillation, with a phase delay between the extracellular field and the transmembrane polarization, as a consequence of the low-pass filtering properties of the neural membrane^139^. If stimulation frequencies are matched to neuronal resonances, effects may accumulate over multiple cycles, and fields as low as 0.2–0.5 V/m have been shown to be capable of shifting spike times *in vitro*^137,140,141^. However, such results should be viewed cautiously when extrapolating to living animals given the irregularity of *in vivo* oscillatory patterns^138^.

Regarding the effects of applied fields on neural membrane dynamics and synaptic noise, it has been observed that interspike intervals (ISIs) in cortical neurons are highly irregular for both evoked^142^ and spontaneous activity^143^. However, *in vitro* experiments in preparations of cortical slices^144^ and the results of detailed simulations^145^ indicate that such apparent stochasticity may instead reflect sensitivity to relatively weak synaptic input fluctuations. Specifically, spike time precision appears to increase for faster stimuli, especially those that recreate synaptic input statistics^144^. This implies that reductions in stimulus time constant and increases in applied field amplitude should lead to increased membrane ramp slope on the depolarizing phase, thus reducing spike time variance attributable to synaptic noise^139^.

While past consensus held that APs were binary, stereotyped events, developments in neurophysiology have presented a more nuanced picture wherein subthreshold polarization of the axonal membrane may induce changes in the AP waveform^146^. These changes may alter the polarization of axon terminals, which in turn modulates neurotransmitter release probability^147^. In particular, Chakraborty et al. demonstrated that weak, depolarizing DCS (producing an E-field of 5 V/m) tends to flatten AP profiles (decreasing amplitude and increasing midwidth) in axon terminals while increasing area under the curve (AUC), and increasing AP conduction velocity in mouse cortical slices^20^. Conversely, hyperpolarization accentuated AP amplitude, decreased midwidth and AUC, and decreased conduction velocity. Changes in the polarization of presynaptic terminals is likely to influence signaling by inducing a stronger/weaker calcium influx, thereby suppressing/potentiating transmitter release and postsynaptic currents (PSCs)^148^. Furthermore, depolarization-induced synaptic current increases have been shown to cause a cascade of effects, ultimately leading to synaptic depression as the pool of presynaptic releasable vesicles is depleted^148^. However, further *in vitro* experiments on the effects of DCS in rodent cortical slices have also revealed that applied fields have the potential to modulate synaptic gain control, even attenuating or reversing the effects of synaptic depression, depending on the polarity of the field^149^. Changes in synaptic efficacy as a consequence of applied fields are unlikely to be related to presynaptic transmitter release kinetics, as DCS had no effect on the time constants for short-term depression or facilitation. Instead, both postsynaptic membrane polarization and the recruitment of additional afferent axons are the putative mechanisms supporting increased synaptic efficacy *in vivo*, implying that DCS may boost cooperativity between synaptic inputs^149^.

Finally, we note the dependence of neural responses to applied fields on the relative orientation of the E-field and the neural morphology (see also *<u>Biophysical Fundamentals of EM-Neuron Interactions</u>* for a discussion of the AF concept). In a study combining multi-scale computational modeling and rat cortical slices, Rahman et al. showed that the field components of relatively weak DCS (8 V/m) produced differential effects on synaptic efficacy^150^. Specifically, they found that radial fields (i.e., those parallel to the somato-dendritic axis) facilitated/inhibited synaptic efficacy via somatic de/hyperpolarization, whereas tangential fields (i.e., those perpendicular to the somato-dendritic axis, and parallel to the cortical surface), exerted pathway-specific effects mediated by axon terminal polarization^150^. Since the direction of terminal polarization depends on the highly variable orientation of the specific afferent pathway with respect to the applied field, tangential fields have little net effect on synaptic efficacy (but preserve pathway-specific effects). In light of the distinct physiological roles played by various synaptic pathways, such polarization is likely to exert significant functional effects, even if net synaptic efficacy remains unchanged^150^. In another *in vitro* study targeting the interrelation between morphology and response to applied fields, Radman et al. found that basal and apical dendrites polarize in opposite directions, and that polarization at distal axon terminals is significantly greater than at the soma^130^, highlighting that the effects of stimulation are not global. These findings underscore the importance of individual morphology, and the relative orientation of electrodes with respect to cortical gyri and sulci in predicting the influence of tES on neural physiology.

##### Delayed Effects

As described above, subthreshold applied fields have the potential to up- or down-regulate ongoing activity by polarizing the neural membrane, and accordingly may incur long-term changes. These include long-term potentiation (LTP), long-term depression (LTD), and metaplastic changes (activity-dependent physiology that regulates neural plasticity)^151^ mediated by the activation of *N*-methyl-D-aspartate (NMDA), group 1 metabotropic glutamate receptors (group 1 mGluRs)^152^, and brain derived neurotrophic factor^153^, among others.

tES has been shown to alter synaptic activity through similar mechanisms as LTP/LTD, including calcium homeostasis^154^ and NMDA receptor recruitment^155^. However, evidence suggests that tES does not induce LTP directly, but rather alters the likelihood that synapses undergo such changes. Indeed, Fritsch et al. showed that DCS in mouse M1 slices enhances LTP, but only in the presence of concurrent presynaptic activity^156^. Furthermore, animal experiments suggest that LTP and LTD are not sufficient to explain the delayed effects of tES, and far-ranging phenomena including gene activation, *de novo* expression of proteins, morphological changes, modified intrinsic firing properties, and glial cell function are also implicated^157^. For example, DC E-fields have been shown to induce cellular migration *in vitro* (i.e., “galvanotaxis”), likely via calcium dynamics-mediated effects^158^, and also evoke changes in the expression of epidermal growth factor receptors, both leading to long-term network alterations^159^. Additionally, immediate early genes are up-regulated in response to activity, and are believed to support the maintenance of LTP^160^. One such gene, zif268, has been shown to be correlated with the duration of LTP^161^, and is increased following tDCS applied to rat hippocampal slices^162^. Finally, we address the effects of tES on metaplasticity mechanisms, which regulate plastic changes and attempt to maintain activity homeostasis by preventing excessive synaptic strengthening or weakening^163^. In particular, preconditioning neural networks by applying stimulation can result in lasting changes that influence the response to subsequent stimulation applied after some interval has elapsed^163^. Such preconditioning activates homeostatic mechanisms that serve to maintain firing rates, for example by increasing postsynaptic receptor density following exogenous inhibition, causing an increase in excitation (i.e., “rebound”) upon disinhibition^163^.

#### Suprathreshold Electrophysiological Impact

##### Spike Initiation

Neuronal activity may also be evoked directly given sufficient field strengths/application of current, though the mechanisms involved are more complex than a simple linear depolarization of the soma to AP threshold. This assertion is undergirded by several lines of evidence, namely: (i) certain cells exhibit spiking for applied fields of both positive and negative polarity, (ii) the inferred membrane potential at E-field-induced firing threshold is not predictive of the actual somatic AP threshold potential, (iii) excitatory postsynaptic potentials as a result of ongoing activity are likely integrated with exogenous E-fields, (iv) DC chronaxie values (minimum time required to evoke an AP for twice rheobase) are lower for intracellular current injection compared to applied E-fields, and (v) certain cells exhibit intrinsic bursting in response to applied fields, but not for somatic current injection^130^. Such bursting behavior could be explained by the propensity of applied E-fields to simultaneously de- and hyperpolarize different neural compartments, leading to back-propagating APs that impinge on hyperpolarized dendritic arbors and initiate dendritic calcium spikes^130,164^. We note here that the fields required to produce these suprathreshold effects are quite large (30–100 V/m), roughly an order of magnitude larger than those typically employed in tES^130^.

Importantly, the site of AP initiation during suprathreshold exogenous E-field stimulation is understood to occur mostly (or exclusively) in axonal arbors^165–168^. This property has historically generated confusion for the interpretation of certain results in the DBS literature, which, for high-frequency stimulation, simultaneously noted decreased (somatic) activity in the stimulated nucleus^169^ and increased presynaptic activity as measured in the efferent target^170^. Modeling studies of DBS have helped to shed light on the apparent conflict, demonstrating that E-field gradients resulting from cathodic stimulation tend to directly hyperpolarize somatic compartments while depolarizing nodes of Ranvier^171^. Furthermore, somatic and axonal compartments may be partially electrically decoupled, leading to AP initiation in the axon, and suppression of transmembrane voltage at the soma. This decoupling is hypothesized to be enhanced by the relative density of inhibitory transsynaptic inputs on the soma and proximal dendrites, which may be activated in response to extracellular axonal stimulation of presynaptic cells. Simultaneous suppression of somatic compartments and activation of axonal compartments would enhance the electrical decoupling observed in experiments^171^.

##### Conduction Blocking

Another mechanism by which E-fields interact with neural physiology is CB, which consists of a reversible and local inhibition of propagating APs in neural tracts, occurring for high-frequency (>2.5 kHz), localized biphasic electrical stimulation (though CB at frequencies <1 kHz has also been observed)^172^. This effect is typically observed in peripheral nerve stimulation using small, implanted electrodes, with the threshold for CB activation decreasing for small electrode-fiber distances and large fiber diameters^173^. While the mechanisms of CB are known to be related to specific ion channel dynamics, the precise mechanisms have still not been fully elucidated^174^. One hypothesis suggests the complete inactivation of sodium channels due to their fast gating dynamics^175^; another proposes constant activation of the potassium channel^176^. CB was found to have different frequency-dependent dynamics for myelinated and unmyelinated fibers^134^, with myelinated fibers showing a monotonic increase of the CB threshold up to 50 kHz^177–179^, and unmyelinated fibers a non-monotonic dependence^179,180^. As thresholds for CB are higher than those for AP generation, it is currently believed that the comparatively low frequencies and current magnitudes (several mA, and ∼100 Hz, respectively) employed in typical tES applications are not sufficient to induce CB. However, CB has been considered a possible mechanism of action during TIS in the proximity of the stimulating electrodes due to the intense E-fields in the kHz range^95^. It must be noted that relatively large currents may be required in TIS to achieve stimulation of very deep structures, or to allow extreme steering of the stimulation locus. Whether CB would affect peripheral nerves in the scalp under such circumstances is not yet known.

#### Non-Electrophysiological Device-Tissue Interaction

Here we summarize the principal physical mechanisms of (adverse) interactions between delivered currents and tissues at the frequencies relevant for brain stimulation. These include phenomena at the electrode-electrolyte interface, electrolyte-tissue interface, and within the tissue, including various electrochemical processes, frequency-dependent current penetration, and thermal effects. Quantities relevant to safety such as the amount of injected charge and the charge density are discussed.

##### Reactions at the Interface

Important electrochemical reactions and biological processes occur at the interface between the electrode and the tissue/electrolyte with which it is in contact. In DBS, a metal electrode directly contacts brain tissue (electrode-tissue interface), while in tES, a conductive gel forms a buffer layer between the electrode and the scalp, giving rise to two boundaries (electrode-electrolyte and electrolyte-tissue). At an electrode-electrolyte interface, charge transfer is mediated by non-Faradaic reactions (no electron transfer across the interface) and Faradaic reactions (electrons move between electrode and electrolyte)^36^. Safe electrostimulation parameters should be identified to avoid the onset of irreversible reactions resulting in damage to the electrode, or the generation of toxic reaction products at the electrode-gel-skin interfaces.

The presence of reversible (e.g., charging and discharging of neural membranes) and irreversible processes at the electrode/electrolyte-tissue interface causes the impedance across the boundary to exhibit strong frequency dependence in response to electrical stimulation, which modifies the effective waveform that reaches the target. One effect commonly observed during long-duration/continuous stimulation is charge accumulation and concomitant electrode polarization due to weak DC leakage currents present in current sources for pulsed or alternating electrical stimulation^181^.

In humans, applied currents evoke a variety of biophysical and physiological changes including alterations in skin conductance, increased blood flow, and sweating, all of which depend on the intensity, frequency, and duration of the current^182^. Changes in skin impedance below tES electrodes are typically measurable within seconds following the delivery of currents^183^. Non-ohmic behavior initiates at relatively small applied DC voltages (>1 V), and skin conductance can fluctuate during stimulation by an order of magnitude or more. Additionally, skin-electrode contact impedance decreases monotonically between 0 and 100 kHz, with the precise relationship depending on the properties of the materials involved. Typically, the quality of the electrode-skin contact is monitored during tES experiments by measuring the contact impedance before and during the application of currents. Changes in the electrode-skin contact (e.g., due to dehydration of the conducting sponges or solidification of the conductive paste) may affect the safe delivery of currents.

##### Thermal Effects

Thermal effects at the frequencies of interest for NIBS and TIS (kHz range) are mainly due to Joule (resistive) heating that results from the motion of ions within the tissues due to voltage differences between the electrodes. Local temperature increases as a result of EM absorption that do not exceed 1°C are considered to be generally safe^84,87^. For medical applications, ()a 2°C increase is considered safe by the FDA^184^ and different international standards, though in special cases (e.g., magnetic resonance imaging; MRI) this limit may be exceeded (particularly under ‘controlled conditions’). The risk of thermal damage is commonly assessed using standardized exposure metrics, such as the volume-averaged SAR, defined as:

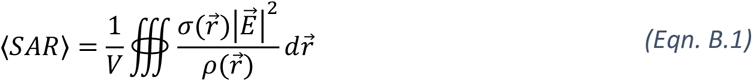

where 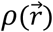 is the mass density. SAR expresses the rate per unit mass at which energy is absorbed by a tissue. However, the exposure-induced heating depends on tissue properties (e.g., perfusion, heat capacity) and the power distribution (which impacts thermal diffusion). Furthermore, thermal sensitivity and the resulting damage depend on many factors, including tissue-specific thermotolerance (i.e., resistance to thermal cytotoxicity), rate of heating^185^, local temperature, and duration of exposure. To estimate the degree of tissue-specific thermal damage, various thermal dose models have been proposed^186^. These include the cumulative equivalent minutes at 43°C dose (CEM43), which converts a transient temperature exposure to an equivalent number of minutes of heat exposure at 43°C in terms of tissue damage. The conversion for periods of near-constant temperature is calculated as

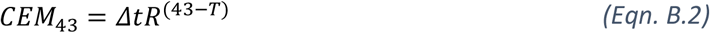

where R is a constant equal to 0.25 for T <43°C and 0.5 for T >43°C (n.b. some references assume *S* = 0 below 39°C). CEM43 safety and/or efficacy thresholds have been proposed for various tissues and applications (e.g., MRI RF safety^187^ and thermal medicine). The lowest thermal dose damage-thresholds have been associated with brain tissue heating and blood brain barrier disruption (2 minutes, while the threshold for skin is 21 minutes according to ISO 14708-2:2019^188^,. CEM43 originates from the Arrhenius tissue damage model, 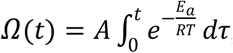, where *e*^−Ω^ represents the surviving cell fraction (or other damage measures), and A and *E*_*a*_ are tissue-specific constants. Unlike the Arrhenius model, CEM43 model shifts the tissue specificity into the damage thresholds, avoiding the use of poorly characterized A and *E*_*a*_ parameters. It also accounts for a known transition in temperature dependence of the damage rate occurring at 42.5–43°C (reflected in the change of the value of R).

It has been argued that temperature increases in the brain >1°C can have long-term effects on brain tissue^189^. While temperature increases in the range of 1°C have been found below tDCS electrodes for 2 mA of current applied for 20 min, the majority of this heating is attributable to the insulating properties of skin, and to changes in blood perfusion as a result of vasodilation (i.e., the “flare effect”)^190^. Current-related heating, in contrast, is known to be orders of magnitude lower. Furthermore, power deposition in brain tissue is considerably lower, and perfusion cooling much more effective, due to extensive vascularization. Antal et al. estimate that power deposition in the brain resulting from 1 mA of applied current is on the order of 0.1 mW/kg, which is about five orders of magnitude less than endogenous metabolic heat production (∼11 W/kg)^17^. In general, the FDA accepts temperature increases of up to 2°C in most tissues (in accordance with some CEM43 definitions that also assume zero contribution to the thermal dose from temperatures below 39°C) and up to 0.1°C in the brain^187,191^.

##### Charge Injection

Damage to tissues associated with the delivery of currents is also dependent on the amount of charge injected within each phase of the stimulation cycle. The charge density is typically compared against the *Shannon limit*^192^, provided by the formula log(*D*) = *k* − log(*Q*) where *D* is the charge density (in μC/cm^2^), *k* is an adjustable parameter (typically between 1.5 and 2), and *Q* is the charge per phase (in μC per phase). This quantity defines the charge threshold at which damage occurs. However, the Shannon limit is chiefly used in the context of bioelectronic medicine where microelectrodes (diameter <300 μm) are common^193^, and is not appropriate for other stimulation scenarios involving large electrode diameters, or for predicting damage far from the electrode. For DBS electrodes, alternative limits for *D* and *Q* were proposed based on tissue damage studies^88,194^.

As an electrode’s charge injection capacity is inversely proportional to its area, the macroelectrodes typical of tES produce charge densities well below those encountered in invasive applications such as DBS and implant-mediated nerve stimulation. Furthermore, the pulse shape of most electrical stimulation applications is designed to be charge balanced (e.g., biphasic symmetric or asymmetric) to avoid the buildup of charge within the tissue. Thus, the low current densities and charge-balanced pulse waveforms typical of tES applications (including TIS) mean that tissue damage directly associated with charge injection is highly unlikely.

#### List of Mechanistic TIS Hypotheses

- In a recent *in vitro* study of carbachol-induced gamma oscillations in rodent hippocampal slices supported by computational network modeling, Esmaeilpour et al. explored the mechanisms underlying the response of deep brain regions to TIS^93^. They propose that sensitivity to TIS is determined by the neural membrane time constant, which, for axonal compartments in particular, approaches the kHz carrier frequency (time constant approx. 1–10 ms). On the other hand, their simulations suggest that selectivity is primarily governed by network adaptation mechanisms with response times shorter than the beat frequency, namely, gamma-aminobutyric acid (GABAB) receptor dynamics, and possibly short-term facilitation and depression, and spike-frequency adaptation via ionic current modulation. Future studies that clarify the role of axonal membranes and network adaptation mechanisms in the response of deep brain tissue to TIS are needed.
- Alternatively, in a simulation study of multi-compartment Hodgkin-Huxley axons, Mirzakhalili et al. argue that neural demodulation of the amplitude-modulated stimulus depends on the capacity of cell membranes to perform signal rectification^95^. They observe that rectification prior to low-pass filtering is a method for demodulating multiplied and convolved signals^195^, and could be achieved in practice through nonlinearities in axonal ion channel dynamics. In particular, simulations revealed that rectification occurred due to differences in the conductance values between sodium and potassium channels, and differences in the speed of activation versus inactivation in sodium channels^95^. These properties were tied to the resonance frequency of the axon and predicted the optimum beat frequency for TIS.
- Similarly, Cao et al. advance the argument that both FitzHugh-Nagumo and Hodgkin-Huxley neuron models demodulate amplitude-modulated signals through transient charge imbalances^75^. As above, fast, depolarizing sodium currents activate with increases in the envelope. Hyperpolarizing potassium channels respond more slowly, leading to transient charge accumulation inside the cell, briefly depolarizing the membrane. Here, the key quantities responsible for sensitivity to TIS are ion channel time constants, in contrast to Mirzakhalili et al., who emphasize the intrinsic membrane time constant of axonal fibers^95^.
- In the same vein, Plovie et al. simulate various single compartment neuron models to explore TI mechanisms^196^. To this end, they define a “TI zone” as the range of input currents over which an amplitude modulated TI signal induces firing at the difference frequency while an unmodulated carrier at the same frequency does not. They test for the presence of a TI zone and, if detected, the associated parameter values, in the following compartment-based models: Hodgkin-Huxley (HH), Frankenhaeuser-Huxley (FH), leaky integrate-and-fire (LIF), exponential integrate-and-fire (EIF), and adaptive and exponential integrate-and-fire (AdEx). Furthermore, they investigate whether nonlinearities in the Goldman-Hodgkin-Katz (GHK) flux equation could be responsible for the neural response to TIS by comparing versions of the FH and HH models in which this term has been linearized (or not) using a first-order Taylor expansion. They find that LIF models (and variants) cannot reproduce experimentally known responses to TIS, while the FH and HH models (and linearized variants) can. Thus, they conclude that GHK nonlinearity is not a necessary driver of the neural response to TIS. Furthermore, by varying the time constants and steady-state values of the FH and HH ion channels, they demonstrate that the response to TIS exposure depends sensitively on nonlinearities in voltage-dependent ion channel gating dynamics^196^.

## Appendix C

### Data Collection

Information regarding AEs and brain stimulation safety were primarily collected from the scientific literature and complemented with clinical trial data, and premarket approval (PMA) and postmarketing surveillance (PMS) data provided by regulatory agencies.

#### Scientific Literature Search

A systematic literature search was performed between February and April 2021, and between September and October 2022, for articles published in peer-reviewed journals using PubMed, Web of Science, Sciencedirect, Google Scholar, Medline and Cochrane Library, for each of the following four technologies: TIS, tACS, tDCS, and DBS. The searches for each technology were conducted by two team members independently (for TIS by a single team member) and included the following search terms:

i. TIS: “TI” + “brain stimulation”, “temporal interference” + “brain”, “temporal interference” + “brain stimulation”, “temporal interference stimulation”.
ii. tACS/ tDCS: “tDCS”, “transcranial direct current stimulation”, “tACS”, “transcranial alternating current stimulation, “tES”, or “transcranial electrical stimulation” in combination with “adverse events”, “side effects”, “adverse effects”, “long term effects”, “neural plasticity”, “frequency-dependent”, “frequency-specific” + “effects”, “safety”, “risks” and with the following adverse event types: “hypomania”, “mania”, “seizure”, “depression”.
iii. DBS: “deep brain stimulation” or “DBS” in combination with “adverse events”, “adverse effects” or “side effects”; also “DBS” or “adverse events”, “adverse effects” or “side effects” in combination with the following disease types: “Parkinson”, “Tremor”, “Dystonia”, “Tourette”, “Obssessive Compulsive Disorder”, “Depression”, “Major depression”, Epilepsy”, “Addiction”, “Anorexia Nervosa”, “Alzheimer”, “Pain”, “Tinnitus”.

For TIS, original research articles concerning experimental results (including simulations) and theoretical explanations for the mechanism(s) of action of TIS were considered. For tACS, tDCS and DBS, we focused on review papers, with a preference for recency; select original articles were also considered as needed to clarify ambiguities identified in the reviews. Publications were scanned for overall quality and relevance with respect to AEs. Only reviews reporting AEs in humans were considered eligible; these were examined in detail to extract specific information on AEs and, where possible, stimulation parameters and target brain structures. For DBS, the type of disease was noted. For tACS and tDCS, we used 10 reviews published between 2011 and 2020, and for DBS a total of 36 reviews published between 2014 and 2021 plus two reviews published in 2003 and 2007, respectively.

#### Regulatory and Clinical Trial Data

The collection of information regarding AEs related to the use of medical devices, for either commercial or investigational purposes, is an obligation for manufacturers and sponsors of clinical investigations. We collected information on AEs from three main sources:

##### PMA Data

PMA data, collected by the FDA, contain information provided by the manufacturer regarding the safety and efficacy of class III medical devices^197^. We extracted information regarding AEs for three prominent commercial DBS devices from available summary reports of device safety and effectiveness.

##### PMS Data

PMS data is information provided by medical device manufacturers, importers, and health care professionals regarding the occurrence of AEs or any other undesirable experiences associated with the use of a regulatory approved medical device in a patient, or voluntary reports by patients and consumers to national regulatory agencies. Such data are publicly available in the FDA’s *Manufacturer and User Facility Device Experience* (MAUDE) data portal^198^. Given the data heterogeneity, we used these reports exclusively to highlight available information, and not to assess the risk profile of any specific medical technologies or products. Searches in the MAUDE portal were conducted within the *Total Product Life Cycle* (TPLC) database using product codes provided partly by a recent publication that lists the FDA product codes of more than 1200 electrical stimulators categorized as tES devices according to their off-label use^199^. Data collected from the MAUDE database cover the period up to April, 2021. We investigated AEs related to electrical stimulation reported in PMS data for the categories *Nervous System, Mental Emotional and Behavioral Disorders, Musculoskeletal System*, and *Skin and Subcutaneous Tissue* for the product codes MHY (Stimulator Electrical, Implanted, for Parkinson Tremor), NHL (Stimulator, Electrical, Implanted, for Parkinsonian Symptoms) and OLM (Deep Brain Stimulator for OCD).

##### EM Safety Guidelines

The ICNIRP 2010^84^ and IEEE Std C95.1-201^87^ safety guidelines, developed to prevent any hazardous effects of exposure to EM fields, were also reviewed. However, these guidelines provide restrictions on unintentional exposure to external electric, magnetic and EM sources, and therefore do not apply directly to an evaluation of TIS safety.

## Footnotes

† The maximal absolute eigenvalue of the Hessian matrix is the directional second derivative along the corresponding eigenvector (*d*^*T*^*Hd* = *λ*_*d*_), which is proprtional to the AF of a fiber oriented in this direction (see *Appendix B*). Here, we report “normalized” AF values by dividing out this scaling factor, which depends on specific properties of the axon in question (i.e., diameter, axoplasmic resistivity, and membrane capacitance).

† ICNIRP guidelines specify that, independent of contact area, currents of ≤1 mA for f ≤2.5 kHz and ≤0.4 x f mA (f in kHz) for f >2.5 and ≤100 kHz are safe for occupational exposure. Thus, extrapolated to a neurostimulation context, field exposures and stimulation currents with applied frequencies above 2.5 kHz would be safer than those of the same amplitude at lower frequencies. However, occupational safety guidelines are of limited applicability to electrical neuromodulation, where stimulation is applied intentionally under controlled conditions. By contrast, safety guidelines provide limits for unintended contact currents and accept non-hazardous sensations as benign side-effects^84^.

